# Machine learning-guided design of artificial microRNAs for targeted gene silencing

**DOI:** 10.64898/2026.02.12.705546

**Authors:** Agnieszka Belter, Jaroslaw Synak, Marta Mackowiak, Anna Kotowska-Zimmer, Marek Figlerowicz, Marta Szachniuk, Marta Olejniczak

**Affiliations:** Institute of Bioorganic Chemistry, Polish Academy of Sciences, Noskowskiego 12/14, 61-704, Poznan, Poland; Faculty of Computer Science and Information Technology, West Pomeranian University of Technology, Zolnierska 49, 71-210, Szczecin, Poland; Institute of Computing Science, Poznan University of Technology, Piotrowo 2, 60-965, Poznan, Poland

**Keywords:** artificial microRNA, RNA interference, amiRNA design, machine learning, gene therapy

## Abstract

Artificial microRNAs (amiRNAs) offer a powerful strategy for targeted gene silencing, but their rational design is limited by complex sequence-structure-processing relationships and the lack of tools capable of optimizing efficacy and specificity. To address this need, we developed miRarchitect, a web-based platform that uses machine learning to support the customizable design of amiRNAs. miRarchitect integrates neural network-guided target-site selection, siRNA insert design, and scaffold choice, utilizing large-scale data from human primary microRNAs (pri-miRNAs) and next-generation sequencing. The platform generates molecules that closely resemble endogenous pri-miRNAs and includes comprehensive off-target analysis to enhance specificity. Experimental validation targeting TMPRSS2 and ACE-2 confirmed precise processing, robust knockdown, and high specificity of miRarchitect-designed amiRNAs. In comparative benchmarking, miRarchitect consistently produced functional amiRNAs, whereas only half of the top candidates generated by other tools showed measurable activity. miRarchitect is freely available at https://rnadrug.ichb.pl/mirarchitect and provides an intuitive interface with an automated workflow for generating, ranking, and selecting candidate amiRNAs for research and therapeutic applications.

## Introduction

RNA interference (RNAi) is a conserved biological process in which small RNA molecules, such as endogenous microRNAs (miRNAs) or small interfering RNAs (siRNAs), regulate gene expression in a sequence-specific manner through interactions with protein partners (Elbashir, Lendeckel and Tuschl 2001; Zeng and Cullen 2003; Bartel 2004). The discovery of miRNAs, in particular, was a landmark that revealed the central role of small RNAs in post-transcriptional gene regulation (Wightman, Ha and Ruvkun 1993; Reinhart *et al*. 2000; Lee *et al*. 2003) – a breakthrough recognized with the 2024 Nobel Prize in Physiology or Medicine awarded to Victor Ambros and Gary Ruvkun (Press release. NobelPrize.org. Nobel Prize Outreach AB 2024. Wed. 7 Oct 2024.). This finding catalyzed the development of RNAi-based technologies for targeted gene silencing (McManus *et al*. 2002; Silva *et al*. 2005). Among these, artificial miRNAs (amiRNAs) have emerged as particularly promising therapeutic agents, offering long-term siRNA expression with reduced off-target effects and cellular toxicity (Silva *et al*. 2005; Castanotto *et al*. 2007; Rao *et al*. 2009).

AmiRNAs are engineered by replacing the endogenous miRNA sequence in a primary miRNA (pri-miRNA) scaffold with a target-specific siRNA. The scaffold ensures proper processing and transport, while the siRNA guides recognition of complementary sequences in target transcripts. Like natural pri-miRNAs, amiRNAs undergo a two-step maturation process: RNase Drosha cleaves the pri-miRNA into a precursor miRNA (pre-miRNA), which is then transported to the cytoplasm by exportin 5 (XPO5) and further processed by Dicer, together with its cofactor TRBP and protein kinase R-activating protein (PACT), into a mature siRNA duplex. Unlike short hairpin RNAs (shRNAs), amiRNAs do not saturate the endogenous miRNA biogenesis machinery, thereby reducing toxicity risks (Castanotto *et al*. 2007; Boudreau, Monteys and Davidson 2008; Amendola *et al*. 2009). Their expression can also be tightly regulated using tissue-specific or inducible promoters, enhancing their therapeutic potential.

Preclinical studies have demonstrated the effectiveness of amiRNAs in targeting disease-associated genes, including those implicated in neurodegenerative disorders – e.g., Huntington’s disease (HD), amyotrophic lateral sclerosis (ALS), neovascular age-related macular degeneration (NAMD) – as well as cancer, cardiovascular disorders, myotonic dystrophies, and viral infections (Borel *et al*. 2016; Pfister *et al*. 2017; Evers *et al*. 2018; Kotowska-Zimmer, Pewinska and Olejniczak 2021; Kotowska-Zimmer *et al*. 2022; Chen *et al*. 2023). The most advanced clinical trial involving amiRNA-based therapy targets the BCL11A gene for the treatment of sickle cell disease (SCD) and is currently in Phase II (NCT05353647). Additional clinical trials are underway at earlier stages for HD (NCT04120493, NCT06826612), oculopharyngeal muscular dystrophy (OPMD, NCT06185673), ALS (NCT06100276), and temporal lobe epilepsy (TLE, NCT06063850). Preliminary clinical data for HD confirm the therapy’s safety and suggest its potential efficacy in slowing disease progression.

Despite this potential, designing effective amiRNAs remains a challenging task. Simply replacing the ∼22-nt miRNA sequence with an exogenous target-specific sequence can change the structure of the resulting amiRNA, disrupt its processing, and ultimately impair its activity. Improperly designed amiRNAs may produce heterogeneous siRNA variants with altered seed sequences or induce arm switching – a phenomenon in which the passenger strand of siRNA is incorporated into the RNA-induced silencing complex (RISC) – thereby increasing the risk of off-target effects (Guda *et al*. 2015; Galka-Marciniak *et al*. 2016; Miniarikova *et al*. 2016). Existing design tools are limited by their reliance on non-human scaffolds (e.g., plant pri-miRNAs (Schwab *et al*. 2006; Ossowski, Schwab and Weigel 2008; Flores-Sandoval *et al*. 2016; Teotia *et al*. 2023)) or a single optimized scaffold (e.g., miR-30). These tools often do not evaluate or optimize the structural integrity of the resulting amiRNA, nor do they adequately assess structural compatibility or off-target risks (Pelossof *et al*. 2017; Park *et al*. 2025).

To address these limitations, we developed miRarchitect, a machine learning-guided web platform for the rational design of human amiRNAs. Leveraging experimentally annotated human pri-miRNA data from MirGeneDB (Clarke *et al*. 2025) and next-generation sequencing (NGS) data, miRarchitect optimizes siRNA insert design, scaffold selection, and structural fidelity while minimizing off-target effects. We validated its utility by designing amiRNAs targeting the TMPRSS2 and ACE-2 transcripts, demonstrating precise processing, robust knockdown, and high specificity in human cell culture. Accessible to non-specialists, miRarchitect enables researchers to generate and rank amiRNA candidates for diverse therapeutic and functional genomics applications.

## Methods

### Architecture and implementation of miRarchitect

The miRarchitect system was built on a two-component architecture consisting of a computational engine (back end) and a web application (front end). The back-end layer, implemented in Python 3.10, integrates key external libraries for RNA data processing, machine learning-based computation, and task management. It utilizes the ViennaRNA package for RNA secondary structure prediction and energy computation (Lorenz *et al*. 2011), TensorFlow 2 (with Keras) for designing, optimizing, and scoring artificial miRNAs via a feedforward neural network (FNN), which serves solely as an accelerator heuristic rather than a predictor of biological silencing efficiency, Flask (with CORS) for handling HTTP requests, and RQ-Scheduler for task scheduling. The back end also relies on an internally constructed database of natural human pri-miRNAs, generated initially from MirGeneDB 2.1 – a manually curated repository containing over 16,000 miRNA genes from 75 metazoan species and more than 1,500 miRNA families (Fromm *et al*. 2015, 2020, 2022) – and subsequently updated using MirGeneDB 3.0 (Clarke *et al*. 2025). Each entry in the miRarchitect database includes: (i) a unique identifier (e.g., hsa-mir-122), (ii) the sequence divided into structural regions (head, prefix, guide/passenger, middle, suffix, passenger/guide, tail), (iii) annotation of the guide strand’s location (5’- or 3’-arm of the pri-miRNA), (iv) the predicted secondary structure, (v) Shannon entropy values for each nucleotide, and (vi) nucleotide-specific free energy values. This curated database serves as the reference source for all computational analyses performed by the miRarchitect engine. The entire system is containerized using Docker to ensure portability and scalability. Its direct installation on a host system has been successfully tested on Ubuntu 22.04.

The front end, developed using React 18 with the Next.js framework, provides an interactive user interface hosted with Nginx. It enables users to submit input data, track computational progress, and retrieve results. The back end continuously updates task statuses – categorized as pending, in progress, finished, or erroneous – to provide real-time feedback. Additionally, the front-end layer connects to the Ensembl database to retrieve extended gene and transcript information (Dyer *et al*. 2025). To enhance usability and accessibility, the interface is designed using Ant Design (5.15.2), ensuring a color-blind-friendly experience with advanced filtering and search functionalities. Data visualization is powered by ECharts (5.5.1), which provides interactive heatmaps for an intuitive representation of results.

All communication between the front end and back end is handled via the HTTP protocol, exclusively using JSON objects. By default, the back end operates on port 5005, with an additional HTTP server (e.g., Apache2) typically used to manage request redirection. miRarchitect is available freely at http://rnadrug.ichb.pl/mirarchitect. It is hosted and maintained by the Institute of Bioorganic Chemistry, Polish Academy of Sciences, Poland.

### The FNN model in miRarchitect

In miRarchitect, we implemented a feedforward neural network as a computational accelerator to pre-screen large numbers of candidate guide-passenger strand combinations before full structural and thermodynamic evaluation. An FNN is an artificial neural network in which information flows strictly in one direction – from the input layer through hidden layers to the output layer – without cycles or feedback loops. Its role in miRarchitect is not to predict biological silencing efficacy, but to approximate the deterministic scoring function and reduce the computational burden of evaluating all possible designs.

The input to the network is the guide-strand sequence, including its prefix, encoded using one-hot encoding. One-hot encoding transforms each nucleotide into a binary vector of length four: A = (1,0,0,0), U = (0,1,0,0), C = (0,0,1,0), and G = (0,0,0,1). Accordingly, a 22-nt guide strand is represented as an 88-dimensional binary input vector. The network produces two outputs: (i) a single continuous value representing the approximated score of the designed miRNA, and (ii) a one-hot encoded pattern specifying which passenger-strand nucleotides remain unpaired.

Because no sufficiently large and diverse experimentally annotated dataset exists that would allow supervised learning of true amiRNA biological activity without overfitting, the FNN has been trained on randomly generated data. Training examples were created by randomly sampling thousands of guide strands and determining the optimal passenger strands limited to nucleotides that are unpaired in the original pri-miRNA secondary structure. For each guide-strand sample, all allowed passenger-strand configurations were evaluated using miRarchitect’s deterministic scoring function, and the configuration yielding the lowest score was selected as the optimal one. All generated examples were retained to ensure robustness across a wide range of sequence configurations.

The model was trained in a supervised manner to learn the mapping from the one-hot encoded guide-strand sequence to its deterministic score and corresponding passenger-strand pattern. The network architecture consists of three hidden layers (128, 256, and 32 units) with SELU activation and a dropout layer (p = 0.2) before the output. Training was performed using the ADAM optimizer (learning rate 0.0003), batch size 256, for 50–150 epochs with MSE loss.

Because the FNN approximates a scoring function derived from experimentally validated principles – including Shannon entropy, nucleotide-resolved free energy, and structural penalties identified in large-scale functional assays of human pri-miRNAs – the synthetic labels used during training remain biologically motivated. Importantly, all designs selected by the FNN undergo a full deterministic re-evaluation within miRarchitect’s pipeline, ensuring that final ranking does not inherit potential FNN bias. Experimental validation is still required to confirm biological activity; however, the FNN substantially accelerates the design process while preserving biological interpretability and determinism of final scoring.

## Results

### Stepwise protocol for ML-guided amiRNA design

The presented framework, implemented in the miRarchitect system, is a fully automatic approach for designing amiRNAs. It analyzes the sequence, structure, and thermodynamics of siRNA duplexes, guide strand/target duplexes, and the final amiRNA as a whole, aiming to construct molecules resembling natural pri-miRNA scaffolds. To efficiently handle the large number of potential candidates and reduce the cost of precise calculations, a neural network is used to preselect candidates for detailed analysis. A key feature of the method is the option of multi-stage off-target filtering, integrated throughout various stages, including target selection, siRNA optimization, and amiRNA construction, minimizing both near-complementary and seed-dependent off-target effects on both strands. The process consists of six steps: (1) input specification, (2) target sequence selection, (3) siRNA design, (4) optimal siRNA selection, (5) amiRNA construction, and (6) scoring and ranking of amiRNA candidates (Figure 1).

**Fig. 1.**
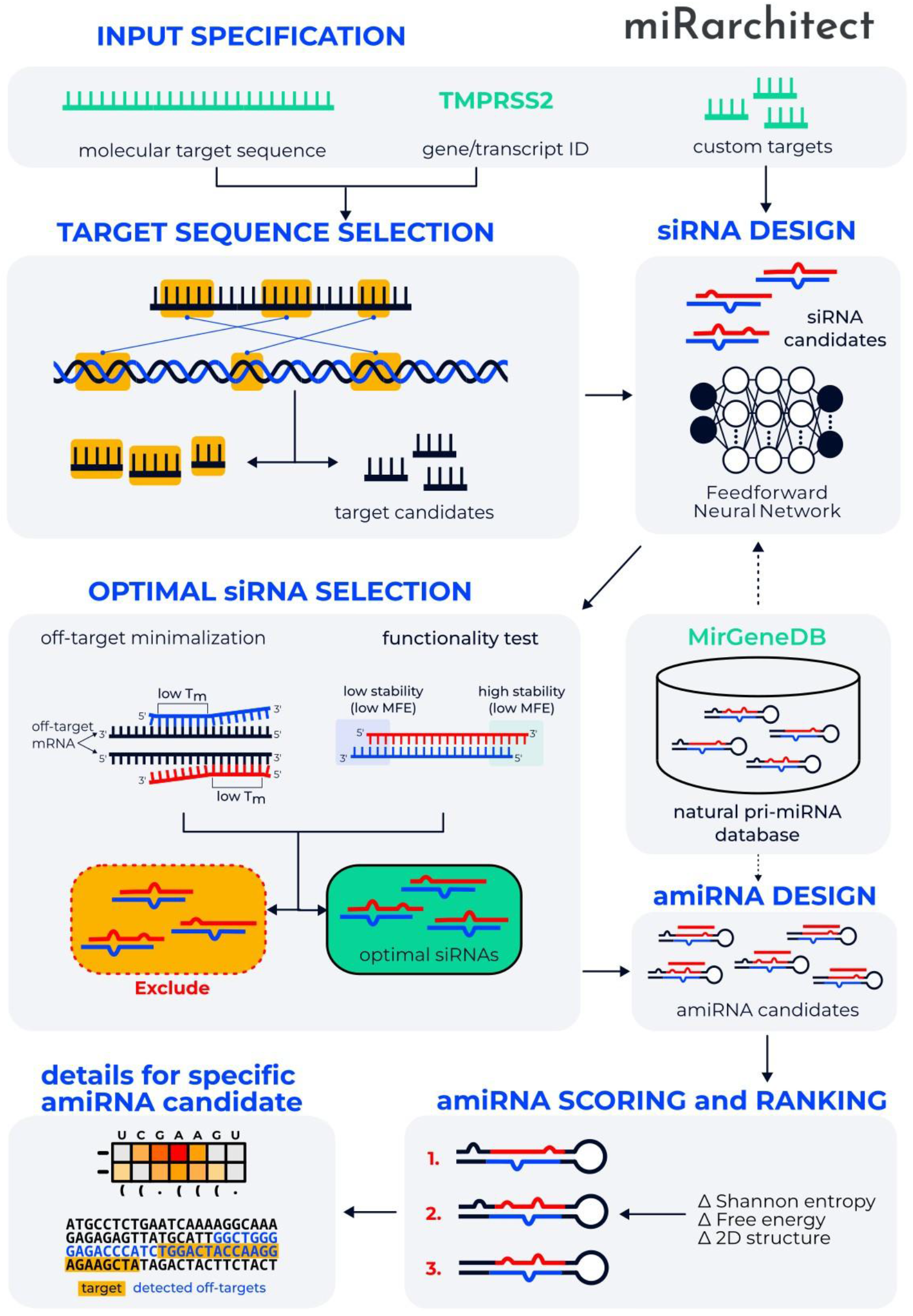
Data flow and processing framework of the miRarchitect system.

The design of amiRNA is guided by the target sequence, which can be provided by the user or identified based on the input data. Both DNA and RNA sequences are accepted. If only an accession number or gene/transcript name is given, miRarchitect generates a list of corresponding transcripts, from which one is selected for processing. The sequence is then analyzed using BLAST, with the longest matching fragment in human DNA/RNA designated as the target, while other alignments are flagged as potential off-targets.

In the second step, the miRarchitect method identifies potential siRNA target sites by analyzing the GC content of each 24-nt fragment of the input sequence. A suitable candidate must avoid GC stretches longer than 9 nucleotides and maintain a GC composition between 30% and 65%. From the pool of preselected targets, only those with minimal near-complementary off-target effects are retained, while sequences with off-targets are discarded. A candidate passes the filter only if it is off-target free. Users can bypass this step, in which case all candidates proceed, and off-targets are merely reported.

The third step involves designing guide and passenger siRNA strands for each previously identified target sequence. miRarchitect selects highly functional siRNAs based on empirically defined rules derived from multiple studies (Amarzguioui and Prydz 2004; Ui-Tei *et al*. 2004). These rules were validated through a high-throughput screening experiment in which more than 71 million human target sequences were tested using the firefly luciferase gene as a reporter (Naito, Saigo and Ui-Tei 2008). The guide strand is fully complementary to its target. The algorithm generates 21-nt RNA duplexes with 2-nt 3’ overhangs (siRNA) using a neural network that provides a preliminary score for each candidate. This score approximates the deterministic scoring function and enables rapid elimination of low-quality designs. Because the FNN is trained on random, self-labelled data rather than experimental ground truth, it serves primarily as an efficient filtering heuristic rather than a source of new biological insight. Its predictions relate to biological viability only indirectly, through the thermodynamic and structural principles encoded in the underlying scoring function. Although the limited sequence space of natural miRNAs constrains the model’s generalizability, the FNN remains a practical and computationally efficient component of the pipeline, with experimental validation ultimately required to confirm biological activity.

To further refine the selection, siRNAs prone to seed-dependent off-target effects are identified and excluded. It is achieved by calculating the melting temperature (Tm) for each molecule. Tm is an essential thermodynamic parameter governing the formation of RNA duplexes, exhibiting a strong positive correlation with the induction of seed-dependent off-target effects, as demonstrated in a large-scale study involving over 16,000 seed sequences (Ui-Tei *et al*. 2008). Therefore, selecting siRNAs with a low Tm for the seed-target duplex is expected to minimize unintended silencing. miRarchitect calculates Tm following the method described in (Freier *et al*. 1986), applying a default threshold of 21.5°C to distinguish nearly off-target-free seed sequences from those more prone to off-target interactions. Users can adjust this threshold if needed. Candidates exceeding the threshold are rejected. Additionally, since off-target effects can arise from both the guide and the passenger strands, siRNAs with sufficiently low seed-target Tm values for both strands are prioritized.

The next step focuses on the evaluation and selection of functional siRNAs. Functional siRNAs are known to exhibit asymmetric stability at the 5’- and 3’-ends. The siRNA guide strand, which contains the thermodynamically less stable 5’-end, is preferentially retained by the RISC (Khvorova, Reynolds and Jayasena 2003; Schwarz *et al*. 2003; Ui-Tei *et al*. 2008). In most cases, the passenger strand of the siRNA loaded onto RISC is cleaved by Ago2 and subsequently degraded (Matranga *et al*. 2005; Rand *et al*. 2005; Leuschner *et al*. 2006). The retained guide strand pairs with the target mRNA (Elbashir, Lendeckel and Tuschl 2001; Hammond, Caudy and Hannon 2001; Martinez *et al*. 2002). To filter out non-functional candidates, miRarchitect analyzes their structures to predict the energy required to unpair both ends of the siRNA duplexes. Two parameters are considered: (i) the number of nucleotides checked on both ends of the duplex to determine if it will start to dissociate from the correct site, and (ii) the minimum free energy difference in kcal/mol between the dissociation energies of both ends of the duplex, with one end requiring lower energy to dissociate first. These parameters are predefined with values of 3 and 2, respectively. Molecules that do not meet these criteria are rejected.

The next stage is the design of amiRNAs. For each siRNA candidate, the method iterates through all pri-miRNAs stored in the miRarchitect database to identify compatible natural pri-miRNA scaffolds, potentially yielding more than one match. For each candidate-scaffold pair, a new molecule is designed to closely resemble the selected natural pri-miRNA in three key parameters: secondary structure, Shannon entropy, and free energy at each nucleotide position of the entire amiRNA. These parameters are known to influence pri-miRNA processing efficiency and precision, and were identified through a large-scale functional assay testing hundreds of primary miRNAs and thousands of single-nucleotide variants (Rice *et al*. 2020).

Finally, the potential of the entire amiRNA molecule is assessed. Each amiRNA candidate is assigned a score, and a final ranking list is generated. miRarchitect calculates the Shannon entropy and binding energy for each nucleotide, along with the secondary structure of the entire amiRNA. The molecules are then compared to their respective natural pri-miRNAs. The sum of squared errors (SSE) for Shannon entropies and binding energies is calculated and added to the final score (Equation 1). If the prefix of the guide strand differs from the natural prefix, a penalty is added to the score. Each difference in the 2D structure between amiRNA and its corresponding natural pri-miRNA also incurs a penalty (an additional 10 points are added to the score). The candidate is rejected if the number of differences in the dot-bracket encoding exceeds the predefined threshold of 4. The remaining designed amiRNAs are displayed in the output and ranked according to their probability of being functional and effective in targeting the selected gene.

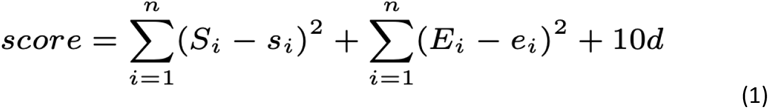

where *n* is the length of amiRNA, *Si* – Shannon entropy of the *i*-th nucleotide in the natural pri-miRNA scaffold, *si* – Shannon entropy of the *i*-th nucleotide in the amiRNA candidate, *Ei* – free energy of the *i*-th nucleotide in the natural pri-miRNA scaffold, *ei* – free energy of the *i*-th nucleotide in the amiRNA candidate, and d is the number of differences in the secondary structure between the amiRNA candidate and the natural pri-miRNA scaffold.

All algorithms used in the miRarchitect’s pipeline — from target site identification, through guide and passenger strand design, functional siRNA selection and scaffold choice, to amiRNA ranking – are grounded in strong scientific evidence derived from experimentally validated principles and high-throughput studies.

### Computational performance and scalability

To evaluate the computational performance of miRarchitect, we conducted runtime benchmarks using synthetic random sequences and biological sequences derived from the BRCA1 transcript, with input lengths ranging from 10 to 5,000 nucleotides. The average runtimes scaled predictably with input size, ranging from 7 seconds for the shortest sequences to 48 minutes for the longest biological sequences under the current web submission limits. Importantly, memory requirements during these computations remained within the capacity of standard workstations and did not constitute a limiting factor. Detailed runtime results are reported in Supplementary Table S1 and Supplementary Figure S1. They confirm the suitability of miRarchitect for large-scale amiRNA design in practice.

### miRarchitect web platform

The described method is implemented in the publicly available miRarchitect web server. It is designed to work with all commonly used web browsers, such as Mozilla Firefox (134.0 and later), Opera (117.0 and later), and Google Chrome (135.0 and later).

The primary input data to miRarchitect is the target sequence, which can be entered directly by the user, retrieved from the Ensembl database (Dyer *et al*. 2025) if an accession number or the name of a gene or transcript is provided instead of a sequence, or uploaded from a FASTA file. Additionally, the web server can be tried for one of six predefined examples available on the homepage. If the user provides a gene or transcript identifier, the system may require further specification by selecting one of the available transcript options. Subsequently, users can choose the pri-miRNA scaffold from a list that contains hsa-mir-21, hsa-mir-30a, hsa-mir-122, hsa-mir-135b, and hsa-mir-155 (hsa-mir-21, -30a, 122, and -155 are selected by default) and adjust amiRNA design parameters. The latter have experimentally determined default values, including the allowed GC content in the target sequence (30-65%), the maximum GC stretch length (9 nts by default), the number of base pairs required to initiate binding (5 bps) and duplex dissociation (3 bps), the minimum difference between the dissociation energies at both duplex ends (2 kcal/mol), the maximum allowed structural variance between amiRNA and the pri-miRNA scaffold (4 nts), and the maximum melting temperature (21.5°C). The users can also decide whether to allow the guide strand to have a different prefix than the pri-miRNA scaffold (enabled by default), activate off-target filtering (disabled by default), and set the invalid prefix penalty, which is added to the amiRNA score if the prefix differs from the natural pri-miRNA (default value: 2.5).

At the output, the web server generates a ranked table of amiRNAs designed by the miRarchitect algorithm, sorted by scores reflecting the predicted functionality and efficiency in knocking down the selected target (Supplementary Figure S2A). The table includes important details for each amiRNA candidate: (i) amiRNA candidate number, (ii) pri-miRNA scaffold, (iii) amiRNA sequence, (iv) target sequence, (v) target start (position of the target site within the input sequence), and (vi) target end (position of the target site endpoint within the input sequence), (vii) score, (viii) end stability difference [kcal/mol] (free energy difference between the dissociation energies of both duplex ends, with one end exhibiting lower energy to be dissociated first), (ix) Tm (melting temperature of the duplex) [°C], (x) minimum off-target mismatches for guide and passenger strand, and (xi) number of off-targets within indicated mismatches. The table is sortable by any numerical parameter, enabling users to organize the data as needed.

Following the main result table, the section on off-target statistics is provided for all amiRNA designs, separately for the guide and passenger strands, and across different regions, including CDS, 5’-UTR, 3’-UTR, and non-coding RNAs (ncRNAs) (Supplementary Figure S2B). It shows how many perfectly matching off-targets or those with at most five mismatches were identified for each candidate. This data enables the filtering of amiRNAs with a high number of off-targets in specific regions, such as the 5’-UTR, or the selection of candidates without off-targets. Upon selecting a specific amiRNA from the main table, further details are provided, divided into several sections. First, the full sequence with annotation is displayed (Supplementary Figure S2C). It highlights the target region(s) complementary to the guide strand and any detected off-target regions. The next section presents structural stability indicators computed based on the comparison of selected amiRNA candidates and the natural pri-miRNA scaffold (Supplementary Figure S2D). Users can view a heatmap showing color-coded values for Δ Shannon entropy and Δ free energy. Beneath the heatmap, the system also displays the secondary structure of the amiRNA candidate encoded in dot-bracket notation. Differences in 2D structure relative to the pri-miRNA scaffold are highlighted for clarity.

Finally, the results page contains a table that shows off-targets with a perfect match and those with up to 5 mismatches (≤17 complementary nucleotides) detected for both the guide and passenger strands (Supplementary Figure S2E). The searches are performed for the 22-nt sequences in both strands of the siRNA duplex, and alignments displayed in the table help identify the mismatch locations. If there are no off-targets with ≤5 mismatches, the table is not shown. For user convenience, all data shown on the results page can be downloaded by clicking the appropriate button under the table of interest. Textual data are saved in CSV format, while graphical data, such as the heatmaps showing Δ Shannon entropy and Δ free energy, are saved in SVG format.

### Experimental validation of the method

The quality of amiRNAs designed by miRarchitect was evaluated in an experiment targeting transmembrane serine protease2 (TMPRSS2) and angiotensin-converting enzyme 2 (ACE-2). These targets were selected because they represent promising candidates for therapeutic intervention against viruses, including influenza A virus (IAV) and severe acute respiratory syndrome coronavirus 2 (SARS-CoV-2) (Batool, Chokkakula and Song 2023). TMPRSS2 mediates the initial step of viral infection through the proteolytic activation of the viral surface glycoprotein, while ACE-2 serves as a major entry receptor, enabling viral penetration into host cells. Inhibition or downregulation of TMPRSS2 and ACE-2 has been shown to prevent the entry of both IAV and SARS-CoV-2 into host cells (Hoffmann *et al*. 2020; Nowak *et al*. 2024).

For the design, human TMPRSS2 variant 1 (TMPRSS2v1; accession NM 005656, gene identifier ENSG00000184012, transcript ENST00000332149) and human ACE-2 (accession NM 021804, a 596-nt consensus sequence corresponding to ACE-2 variants 1–6, coordinates 1304–1900) were used as input. Pri-miRNA-122, -135b, and -155 were chosen as scaffolds (Supplementary Table S2), and advanced miRarchitect parameters were kept at their default settings.

From the ranking list of TMPRSS2-targeting amiRNAs generated by miRarchitect, we selected and compared amiRNAs with low vs. high scores based on the same scaffold (i), and amiRNAs targeting the same sequence but derived from different scaffolds (ii) (Supplementary Figure S3A–B; Supplementary Table S3). Then, the silencing efficiency of the amiRNAs was analysed in a cellular model. HEK293T cells were co-transfected with a vector expressing TMPRSS2 v.1, as well as Renilla and firefly luciferase genes, and pcDNA3.1 plasmid encoding amiRNA. A plasmid encoding natural pri-miRNA was used as a control. Luciferase assay was performed 24 hours after transfection. The results of this experiment confirmed the predictions from miRarchitect scoring, showing that amiRNAs with lower scores were more effective in TMPRSS2 silencing (Supplementary Figure S3C). Moreover, amiRNAs composed of the same siRNA insert and different scaffolds exhibited different silencing efficiencies. The most effective reagents (miR_122_02_T, miR_135b_01_T, and miR_135b_02_T) reduced the TMPRSS2 level by approximately 80% (Supplementary Figure S3C; Supplementary Table S4).

amiRNA activity was also assessed at the transcript and protein levels in a separate experimental setup. HEK293T cells were co-transfected with plasmids encoding the target gene (pcDNA3.1 TMPRSS2/V5) and selected amiRNA (pcDNA3.1 amiRNA). The most significant reduction in TMPRSS2 transcript and protein levels (∼80% and ∼95%, respectively) was observed in cells treated with miR_122_02_T (Supplementary Figures S3–E and S4; Supplementary Tables S5 and S6). Six of the eight tested amiRNAs reduced protein levels by ≥75% (Supplementary Figure S3E).

Similarly, ACE-2-targeting amiRNAs were designed. Two amiRNAs with the lowest (best) scores – one based on pri-miR-122 and the other on pri-miR-155 – were selected from the ranking list (Supplementary Table S3). Both showed comparable, moderate silencing efficiency, reducing ACE-2 protein and transcript levels by approximately 50% (Supplementary Figure S5; Supplementary Tables S7 and S8).

We next analyzed the products of amiRNA processing by Drosha and Dicer in HEK293T cells using small RNA sequencing. As expected, the amount of short RNAs released from most amiRNAs was low, representing <1% of the endogenous miRNA pool for 8 of 10 amiRNAs (Supplementary Table S9). No straightforward relationship was observed between guide strand abundance and TMPRSS2 or ACE-2 silencing efficiency (Supplementary Figures S3 and S5). According to MirGeneDB 3.0, the selected pri-miRNAs exhibit a high guide-to-passenger strand ratio. We confirmed it for pri-miRNA-122 and -155, where 99.6% and 99.8% of products were derived from the 5’-arm (Supplementary Figures S6A and S6B). Interestingly, pri-miRNA-135b released similar numbers of products from both arms. Likewise, pri-miR-135b-based amiRNAs with low scores (high similarity) also released products from both arms (Supplementary Figure S6C).

Nine of the ten designed amiRNAs showed high 5’-end homogeneity for Drosha products. In contrast, ∼15% of products from miR_122_01_T showed a 5’-end shift despite maintaining all parameters similar to the natural scaffold, with structural analysis providing no explanation of this effect (Supplementary Figure S7). Comparison of three amiRNAs with the same siRNA insert but different scaffolds revealed variations in released guide strands, all with uniform 5’-ends but differing in length (22 nucleotides for the most active miR_122_02_T to 24 nucleotides for the least active miR_155_01_T; Supplementary Figure S6). These lengths corresponded to products released from their natural precursors. All remaining experimental details are provided in the Supplementary file.

### Comparison with existing amiRNA design tools

miRarchitect introduces a unique capability for amiRNA design by allowing users to select among multiple human endogenous pri-miRNA scaffolds – a flexibility not offered by other available tools (Supplementary Table S10). Using endogenous scaffolds ensures structural compatibility with the natural miRNA biogenesis pathway, supporting efficient Drosha and Dicer processing. The platform currently provides five well-characterized pri-miRNAs – pri-miR-21, -30a, -122, -135b, and -155 – selected based on validated and precise cleavage patterns documented in MirGeneDB and their successful use in numerous in vitro and in vivo studies (Silva *et al*. 2005; Fellmann *et al*. 2013; Galka-Marciniak *et al*. 2016; Pfister *et al*. 2017; Kern *et al*. 2020; Kotowska-Zimmer *et al*. 2022; http://www.mirnainfo.com/miRNADesigner.aspx). In this work, we focused on scaffolds that release the mature miRNA from the 5’ arm, as the Drosha cleavage at this position precisely defines the seed region essential for target recognition (Lee *et al*. 2003; Silva *et al*. 2005). Scaffolds releasing the product from the 5’ arm, therefore, provide greater reliability for amiRNA design. Among the tested scaffolds, pri-miR-122 and pri-miR-155 produced a single, well-defined siRNA from the 5’ arm, whereas pri-miR-135b generated products from both arms. Due to discrepancies between predicted and observed processing of pri-miR-135b, we do not recommend this scaffold as a first choice. This emphasis on 5’-arm–derived guide strands further distinguishes miRarchitect from tools such as shRNAI and SplashRNA, which predominantly release the guide strand from the 3’ arm.

Other web-based tools developed for amiRNA design include shRNAI (Park *et al*. 2025), SplashRNA (Pelossof *et al*. 2017), miRNA Designer (http://www.mirnainfo.com/miRNADesigner.aspx), WMD3 - Web MicroRNA Designer (Schwab *et al*. 2006; Ossowski, Schwab and Weigel 2008), and amiRNA Design Helper (Flores-Sandoval *et al*. 2016) (Supplementary Table S10). Three of them use scaffold sequences from Homo sapiens and other species, such as *Mus musculus* or *Arabidopsis thaliana*. All but one implement a ranking system to help users identify the most promising amiRNA candidates. However, only miRarchitect and amiRNA Design Helper include an off-target identification module, a crucial feature for minimizing unintended gene silencing.

Among these tools, shRNAI shares the most methodological similarities with miRarchitect. Both employ neural networks for predicting amiRNA effectiveness, although shRNAI utilizes a convolutional neural network (CNN) with three convolutional layers and one regression layer, whereas miRarchitect applies a simpler feedforward neural network. Additionally, while shRNAI predicts only the effectiveness of the guide strand, miRarchitect designs both the guide and passenger strands. Another key difference is that in shRNAI, the CNN model itself provides the final scoring for designed molecules, whereas the miRarchitect’s FNN is integrated into a broader computational framework that includes thermodynamic calculations.

A fundamentally different approach is implemented in SplashRNA. This tool focuses on designing shRNA rather than amiRNA and employs two support vector machine (SVM)-based models trained on experimental datasets. Unlike miRarchitect, it does not incorporate thermodynamic calculations, treating sequences as simple text strings for machine learning-based prediction.

To enable a more meaningful comparison between miRarchitect and existing amiRNA design tools, we carried out a systematic evaluation of their computational behaviour. In practice, however, establishing a unified benchmark proved feasible only to a limited extent due to substantial differences in input formats, algorithmic workflows, and output modes across tools. Among the available platforms, only amiRNA Design Helper allowed reproducible, interpretable runtime measurements for full-length input sequences: using both biological and random RNA transcripts ranging from 100 to 5,000 nucleotides, we observed computing time increased steadily with sequence length, from approximately 9 seconds for 100-nt inputs to 17–23 seconds for 5,000-nt sequences. In contrast, shRNAI accepts sequences only up to ∼3,000 and completes analyses in fractions of a second, preventing meaningful runtime evaluation for longer transcripts. SplashRNA and miRNA Designer require users to provide predefined target sites instead of scanning whole transcripts, making them incompatible with transcript-level benchmarking. WMD3 provides results exclusively via asynchronous email notifications without reporting computation times, with queue-related delays further obscuring algorithmic performance. Despite these limitations, our evaluation indicates that miRarchitect is among the few tools capable of processing long human transcripts fully autonomously – from target-site selection to complete amiRNA design – within practical and reproducible computation times.

To further assess miRarchitect relative to existing design tools, we generated TMPRSS2-targeting amiRNAs using shRNAI and SplashRNA and experimentally evaluated their silencing efficiency in HEK293T cells. Among the six top-ranked molecules produced by these tools, only half induced statistically significant reduction in TMPRSS2 transcript (Supplementary Figure S8A) and protein levels (Supplementary Figure S8B). Notably, highest-scoring candidates predicted by shRNAI and SplashRNA failed to measurably silence TMPRSS2, underscoring the need to test multiple designs to identify functionally active molecules. In contrast, all amiRNAs designed using miRarchitect exhibited strong silencing activity (≥50%), including those assigned less favorable scores (Supplementary Figure S3).

## Conclusion

Post-transcriptional inhibition of gene expression remains a cornerstone of both basic research and therapeutic development. Among the tools enabling this, artificial microRNAs stand out for their ability to harness the endogenous RNAi machinery for sustained gene silencing – a critical advantage over transient approaches like synthetic siRNAs or antisense oligonucleotides (ASOs), which require repeated administration. However, the full potential of amiRNA technology has been limited by challenges in designing molecules that balance structural fidelity, processing efficiency, and specificity. While plant biology has benefited from dedicated amiRNA design tools (Schwab *et al*. 2006; Ossowski, Schwab and Weigel 2008; Flores-Sandoval *et al*. 2016; Teotia *et al*. 2023), human applications have lagged, relying primarily on modified miR-30 scaffolds (Pelossof *et al*. 2017; Park *et al*. 2025) that lack the diversity needed for optimal targeting. miRarchitect addresses this gap by integrating machine learning with a comprehensive human pri-miRNA scaffold library, enabling the design of amiRNAs that preserve native structural features while minimizing off-target effects. Our FNN-driven pipeline automates siRNA insert optimization and scaffold matching, ensuring precise pri-miRNA processing and homogeneous amiRNA maturation. This computational strategy was validated in a case study targeting TMPRSS2 and ACE-2, where miRarchitect-designed amiRNAs exhibited processing profiles indistinguishable from natural pri-miRNAs and achieved robust gene silencing with minimal off-target activity. By incorporating NGS data and structural constraints, miRarchitect outperforms existing tools, which often neglect critical factors such as scaffold integrity and siRNA seed sequence conservation.

The platform’s modular architecture allows for continuous expansion, with new pri-miRNA scaffolds added as MirGeneDB and experimental datasets grow. For example, discrepancies in pri-miRNA-135b processing annotations between MirGeneDB and miRSwitch (Kern *et al*. 2020) underscore the importance of iterative updates to refine scaffold libraries. Furthermore, emerging evidence suggests that regulatory elements in pri-miRNA flanking regions and loops could enable tissue-specific amiRNA expression – a feature we aim to integrate into future versions of miRarchitect.

Despite its advancements, miRarchitect’s predictions require experimental validation due to the context-dependent nature of miRNA biogenesis, including factors such as tissue-specific arm switching and competing endogenous RNA (ceRNA) networks that may influence amiRNA efficacy in vivo. While current off-target analysis helps mitigate sequence-based risks, epigenetic factors and 3D structural interactions remain challenging to predict. A limitation of this study is that experimental validation was performed for only two target genes (TMPRSS2 and ACE-2), which may not fully capture sequence- or context-dependent determinants of amiRNA performance. Moreover, FNN was trained exclusively on random data to approximate the deterministic scoring function, because no sufficiently large experimental dataset exists to support supervised learning of true biological activity. Therefore, experimental validation across additional targets remains essential. Although miRarchitect’s scoring function integrates diverse computational and structural features grounded in established principles, certain biological variables – such as co-factor interactions and tissue-specific regulatory environments – are not yet fully represented. To address these limitations, future work should focus on expanding experimental testing to additional targets and biological contexts, as well as enhancing miRarchitect with more comprehensive off-target reporting and transcriptome-wide visualization. Such improvements will increase user confidence in the safety and specificity of designed amiRNAs and support the development of next-generation RNAi-based therapeutics.

In conclusion, miRarchitect bridges a critical gap in RNAi therapeutics by providing a scalable, user-friendly platform for designing human amiRNAs with precision and specificity. Its ability to balance sensitivity and specificity, coupled with adaptability to evolving genomic data, positions it as a valuable resource for advancing both functional genomics and RNA-based drug development.

## Availability of data and materials

The miRarchitect tool is publicly available at http://rnadrug.ichb.pl/mirarchitect. The small RNA sequencing data have been deposited in the NCBI Sequence Read Archive (SRA) under accession numbers PRJNA1264680 and PRJNA1371627, accessible via the following read-only links: https://dataview.ncbi.nlm.nih.gov/object/PRJNA1264680?reviewer=pq0u3q0ir0pfp7jai80qii5gbc and https://dataview.ncbi.nlm.nih.gov/object/PRJNA1371627?reviewer=4s5fg4igje5faatf49lqfsijst

## Author contributions statement

M.O., M.S., A.B., and J.S. conceived the research. A.B. and J.S. developed an amiRNA design protocol. M.S. and M.M. designed the web application. J.S. and M.M. implemented components of the miRarchitect system. A.B., J.S., M.M., and M.S. tested miRarchitect and performed the experiments. A.B., J.S., M.O., and M.S. analyzed and interpreted the results. A.B., J.S., and M.M. prepared the figures. A.B., J.S., M.O., M.S., A.K.Z., and M.F. drafted, edited, and revised the manuscript. All authors reviewed and approved the final version of the manuscript. M.F. obtained funding and supervised the project.

## Acknowledgments

We thank Dr. Qingyu Wu and Dr. Ningzheng Dong from Soochow University for providing the pcDNA3.1/V5 TMPRSS2v1 plasmid. We also acknowledge Joanna Sarzynska, Agnieszka Rybarczyk, and Michal Smuszkiewicz for helpful discussions on the concept of the miRarchitect tool.

