## Supplemental Data for "Machine learning-guided design of artificial microRNAs for targeted gene silencing"

### Supplementary Information

#### Experimental procedures for validation of amiRNA design

#### Cell culture

HEK293T cells were maintained in Dulbecco's Modified Eagle's Medium (DMEM; Gibco) supplemented with 10% fetal bovine serum (FBS; BioWest), 1% Penicillin-Streptomycin (PenStrep; Gibco), and 2mM L-glutamine (Gibco). Cells were cultured at 37°C in a humidified atmosphere containing 5% $CO_2$. The cell line was routinely monitored for mycoplasma contamination using the Venor→ GeM Classic Mycoplasma Detection Kit for conventional PCR (Minerva Biolabs).

#### Target and amiRNA expression plasmids

The target expression plasmid, psiCHECK2_TMPRSS2v1, contains two reporters: *Renilla* and firefly luciferase genes, under SV40 and HSV-TK promoters, respectively (Promega). The TMPRSS2v1 gene was cloned into XhoI/NotI cloning sites under the control of the SV40 promoter, following the *Renilla luciferase* gene. For the generation of pcDNA3.1_TMPRSS2/V5 plasmid – complementary DNA (cDNA) (NM_001135099.1), encoding human TMPRSS2 (isoform 1), was amplified from human MTC II cDNA panel (Clontech; catalog no. 636743) and inserted into pcDNA3.1/V5 plasmid (Thermo Fisher Scientific; catalog no. K4800-01) (Zhang *et al.* 2020). The target expression plasmid pcDNA3.1-hACE-2 (Addgene; plasmid #145033), with human ACE-2 ectodomain (residues 1-615) (GenBank accession number NM_021804) and a C-terminal C9 tag under CMV promoter, was prepared as described previously (Shang *et al.* 2020).
AmiRNA expression plasmids contain the amiRNA gene under the control of the CMV promoter. The amiRNA gene was synthesized (GeneScript) and cloned into the pcDNA3.1/Hygro(+) plasmid into the HindIII/XhoI cloning site (Thermo Fisher Scientific).

#### Luciferase assay

HEK293T cells ($2x10^4$ cells/well) were seeded into 96-well plates and, after 24 hours, the cells were co-transfected with psiCHECK-2_TMPRSS2v.1 plasmid and pcDNA3.1_amiRNA plasmid in a 1:10 ratio using Lipofectamine 2000 (Invitrogen, Thermo Fisher Scientific, Carlsbad, CA). pcDNA3.1_pri-miRNA plasmids coding natural pri-miRNAs served as controls. Twenty-four hours after transfection, cells were harvested and lysed using Dual-Glo® Luciferase Assay Reagent (Promega, Madison, WI, USA). The bioluminescence assay was performed using a Dual-Glo® Luciferase Assay protocol (Promega) and Victor Nivo Plate Reader (PerkinElmer, Waltham, MA) according to the manufacturer's instructions. The luminescence intensity of firefly luciferase was normalized to the luminescence intensity of *Renilla luciferase*. The efficiency of amiRNAs was determined as the ratio of *Renilla* to firefly luminescence for the amiRNA, relative to the corresponding ratio for the control.

#### RT-qPCR analysis

HEK293T cells ($9x10^5$ cells) were seeded into a 3.5 cm dish and, after 24 hours, the cells were transfected with 4 µg of pcDNA3.1_amiRNA using 6 µg Lipofectamine 2000. After 2 hours, the cells were transfected with 2 µg of pcDNA3.1_TMPRSS2/V5 or pcDNA3.1- ACE-2 plasmid using 6 µl Lipofectamine 2000. Twenty-two hours after the second transfection, the cells were collected, and RNA was isolated using Total RNA Mini Plus D (AA Biotechnology) according to the manufacturer's instructions. A DeNovix Nanodrop Spectrophotometer was used to measure the RNA concentration. RT-qPCR was performed in the CFX Connect Real-Time PCR Detection System (Bio-Rad, Hercules, CA) in a 10 µl total volume of reaction, using 50 or 100 ng of total RNA. 200 nM of each forward and reverse TMPRSS2 primers or 120 nM of each forward and reverse ACE-2, with β-actin and GAPDH as the reference gene with 150 nM concentrations of relevant forward and reverse primers, using iTaq™ Universal SYBR® Green One-Step Kit (Bio-Rad), under the following conditions: reverse transcription reaction at 50°C for 10 min, denaturation at 95°C for 30 s followed by 40 cycles of denaturation at 95°C for 15 s and annealing at 60°C for 30 s. Sequences of specific primers are listed in Supplementary Table S11.

**Protein extraction and Western blotting**

Cell lysis and protein extraction were done using RIPA buffer (Thermo Scientific) according to the manufacturer's protocol. 20 µg of total protein per sample was mixed with 4x Laemmli sample buffer (Bio-Rad) containing 5% β-mercaptoethanol, heated at 95°C for 5 min. The sample was loaded into the well of SDS-polyacrylamide 4-15% Mini PROTEAN® TGX™ (Tris-Glycine-eXtended) Precast Gels (Bio-Rad). The electrophoresis was performed in Tris/Glycine/SDS running buffer (Bio-Rad) at 150 V. Then, the proteins were transferred to a nitrocellulose membrane (Amersham Protran 0.45 NC. Cytivia) by the wet transfer method in Tris/glycine/methanol buffer. at 4°C, 100 V, for 1h. The membrane was blocked with 10% milk in PBS/0.1% Tween 20 buffer. The primary and secondary antibodies were used in PBS/0.1% Tween 20 buffer containing 3% BSA. Immunoreactions were detected using Westar Supernova (Cyagen, XLS3.0100). Protein band densities were quantified using a Gel-Pro Analyzer (Media Cybernetics). Antibody details are provided in Supplementary Table S12.

**Analysis of amiRNA processing by small RNA sequencing**

HEK293T cells ($9 \times 10^5$ cells/6 cm dish) were transfected 24 hours post-seeding with 4 µg of pcDNA3.1_amiRNA plasmid using 12 µl Lipofectamine 2000 (Invitrogen, Thermo Fisher Scientific, Carlsbad, CA) according to the manufacturer's protocol. Total RNA was isolated 24 hours post-transfection using the miRVanaTM miRNA Isolation Kit (Thermo Fisher Scientific, Vilnius, Lithuania). RNA integrity was assessed on a Tape Station 4150 (Agilent Technologies, Santa Clara, CA). Small RNA sequencing was performed by Novogene (Cambridge, UK) using an Illumina SE50 sequencing with 10M base pair reads. Adapter-trimmed reads were quality-filtered and converted to FASTA format using a custom Python script. Reads were mapped to amiRNA sequences using Bowtie with parameters '-v 0' (no mismatches) and '-a' (report all alignments) (Langmead *et al.* 2009). Mapped reads were visualized in HTML format using a custom Python script for manual inspection of processing fidelity. For miRNA quantification, mature miRNA sequences were retrieved from miRbase (Kozomara, Birgaoanu and Griffiths-Jones 2019), and read counts were normalized to reads per million (RPM) using a separate in-house Python pipeline.

**Analysis of the amiRNA and pri-miRNA structure**

Secondary structures of designed amiRNAs and their corresponding pri-miRNA scaffolds were predicted using RNAfold (Gruber *et al.* 2008; Lorenz *et al.* 2011). These predictions were used as input for RNAComposer (Popenda *et al.* 2012) with default parameters to generate 3D structure models. amiRNA models and their corresponding pri-miRNA scaffolds were energy-minimized and superimposed in PyMOL (Schrödinger, LLC (2016) The PyMOL Molecular Graphics System). The PyMOL Molecular Graphics System) for structural comparison. Root mean square deviation (RMSD) values were calculated for full-length structures and key functional regions. Differences in conformation were quantified to assess structural fidelity between designed amiRNAs and their native pri-miRNA scaffolds.

**Statistical analysis**

All experiments were performed at least three times, each in a minimum of three technical replicates. Comparisons between the two groups were analyzed using an unpaired t-test. For comparisons involving more than two groups, one-way ANOVA was applied, followed by Tukey's HSD test for post hoc analysis. A p-value $< 0.05$ was considered statistically significant. Correlations were assessed using Pearson's correlation coefficient (r). Exact p-values are indicated on the graphs (Supplementary Figures S3, S5 and S8).

**Supplementary Table S1.** The miRarchitect's scalability for large-scale inputs. A comprehensive benchmark was conducted using both biological sequences (fragments of the BRCA1 transcript) and random sequences, with input sizes ranging from 10 to 5,000 nucleotides, reflecting the upper limit currently allowed by the submission webpage. For each input length and category, six independent runs with freshly generated sequences were performed to ensure reliability, and the average runtime for each scenario was calculated. Memory consumption did not exceed standard workstation capacities and did not present a bottleneck under the tested conditions. A graphical representation of the same data is provided in Supplementary Figure S1.

| Sequence length [nt] | Average computing time [s] | |
|---|---|---|
| | Biological sequences | Random sequences |
| 10 | 7.67 | 7.00 |
| 100 | 144.50 | 50.33 |
| 200 | 275.67 | 65.17 |
| 500 | 566.83 | 221.00 |
| 1,000 | 1,105.67 | 350.83 |
| 2,000 | 1,985.17 | 896.17 |
| 5,000 | 2,847.33 | 1,212.33 |

**Supplementary Table S2.** Natural pri-miRNA scaffolds implemented in miRarchitect. Pri-miR-21, -30a, -122, and -155 are selected by default during amiRNA design. The mature miRNA strand is shown in red, and the miRNA* strand is shown in blue.

| **Scaffold** | **Sequence (5'-3')** |
|---|---|
| Pri-miR-21 | ACAUCUCCAUGGCUGUACCACCUUGUCGGGUAGCUUAUCAGACUGAUGUUGACUGUUGAAUCUCAUGGCAACACCAGUCGAUGGGCUGUCUGACAUUUUGGUAUCUUUCAUCUGACCAUC |
| Pri-miR-30a | GAAGGUAUAUUGCUGUUGACAGUGAGCGACUGUAAACAUCCUCGACUGGAAGCUGUGAAGCCACAGAUGGGCUUUCAGUCGGAUGUUUGCAGCUGCCUACUGCCUCGGACUUCAAGGGGCUAC |
| Pri-miR-122 | CGUGGCUACAGAGUUUCCUUAGCAGAGCUGUGGAGUGUGACAAUGGUGUUUGUGUCUAAACUAUCAAACGCCAUUAUCACACUAAAUAGCUACUGCUAGGCAAUCCUUCCCUCGAUAA |
| Pri-miR-135b* | GCCGCUGGACCCCUCCACUCUGCUGUGGCCUAUGGCUUUUCAUUCCUAUGUGAUUGCUGUCCCAAACUCAUGUAGGGCUAAAAGCCAUGGGCUACAGUGAGGGGCGAGCUCCUUCUCCUGC |
| Pri-miR-155 | UGCUGAAGGCUUGCUGUAGGCUGUAUGCUGUUAAUGCUAAUCGUGAUAGGGGUUUUUGCCUCCAACUGACUCCUACAUAUUAGCAUUAACAGUGUAUGAUGCCUGUUACUAGCAUUCACAU |

* The pri-miR-135b scaffold, which may generate mature miRNAs from both arms, is not selected by default, as such dual-arm processing is generally not recommended for therapeutic applications.

**Supplementary Table S3.** Artificial miRNAs targeting TMPRSS2 and ACE-2, designed using miRarchitect, shRNAI, and SplashRNA, and experimentally tested in this study. Guide strands are highlighted in red, whereas passenger strands are highlighted in blue.

| Name | Sequence (5'-3') | Score* |
|---|---|---|
| miRarchitect-designed amiRNAs | | |
| Targeting TMPRSS2 | | |
| miR_122_01_T | CGUGGCUACAGAGUUUCCUUAGCAGAGCUGUUCAUAGACAUAUCUGCUGUUGUGUCUAAACUAUCAACAGCAGAUCUGUCUAUUUACAGCUACUGCUAGGCAAUCCUUCCCUCGAUAA | 5.61 |
| miR_122_02_T | CGUGGCUACAGAGUUUCCUUAGCAGAGCUGUUUAUAGAUAUCGACAUUGCCGUGUCUAAACUAUCAGCAAUGUCGCUAUCUAUGCACAGCUACUGCUAGGCAAUCCUUCCCUCGAUAA | 57.41 |
| miR_122_03_T | CGUGGCUACAGAGUUUCCUUAGCAGAGCUGUACUAUUCCUUGGCUAGAGUAAUGUCUAAACUAUCAACUCUAGCCUAGGAAUAACACAGCUACUGCUAGGCAAUCCUUCCCUCGAUAA | 98.2 |
| miR_155_01_T | UGCUGAAGGCUUGCUGUAGGCUGUAUGCUGUUUAUAGAUAUCGACAUUGCCGUUUUUGCCUCCAACUGACGGCAAGUCAUAUCUAUAAACAGUGUAUGAUGCCUGUUACUAGCAUUCACAU | 4.34 |
| miR_155_02_T | UGCUGAAGGCUUGCUGUAGGCUGUAUGCUGUCUCAAAAUCCCCGCAAAUGCCUUUUUGCCUCCAACUGAGGCAUUGCGGGAUUUUGAGACAGUGUAUGAUGCCUGUUACUAGCAUUCACAU | 102.06 |
| miR_135b_01_T | GCCGCUGGACCCCUCCACUCUGCUGUGGCCUUCAGUUUCAUAAAGCUGGUGGAUUGCUGUCCCAAACUCCACCAGCUAUUGAAACUGAAGGCUACAGUGAGGGGCGAGCUCCUUCUCCUGC | 11.37 |
| miR_135b_02_T | GCCGCUGGACCCCUCCACUCUGCUGUGGCCUUUAUAGAUAUCGACAUUGCCGAUUGCUGUCCCAAACUCGGCAAUGUGUUAUCUAUAAAGGCUACAGUGAGGGGCGAGCUCCUUCUCCUGC | 44.19 |
| miR_135b_03_T | GCCGCUGGACCCCUCCACUCUGCUGUGGCCUCCAAUAGCUGGUGGUGACCCUAUUGCUGUCCCAAACUCGGGUCACCUAAGCUAUUGGAGGCUACAGUGAGGGGCGAGCUCCUUCUCCUGC | 96.44 |
| Targeting ACE-2 | | |
| miR_122_01_A | CGUGGCUACAGAGUUUCCUUAGCAGAGCUGUGGAAUUGGUAAAGGGUCCUUGUGUCUAAACUAUCAAGGACCCUUAACCAAUUAAACAGCUACUGCUAGGCAAUCCUUCCCUCGAUAA | 6.81 |
| miR_155_01_A | UGCUGAAGGCUUGCUGUAGGCUGUAUGCUGUUUAAAUGCUUAGGUGUGGCUGUUUUUGCCUCCAACUGACAGCCAACCAAGCAUUUAAACAGUGUAUGAUGCCUGUUACUAGCAUUCACAU | 11.04 |
| shRNAI-designed amiRNAs | | |
| Targeting TMPRSS2 | | |
| miR-E_01_T | UGCUGUUGACAGUGAGCGACACCAGGAAGGUCAUAUUGAAUAGUGAAGCCACAGAUGUAUUCAAUAUGACCUUCCUGGUGGUGCCUACUGCCUCGGA | 0.7436 |
| miR-E_02_T | UGCUGUUGACAGUGAGCGAGACUACCAAGGAGAAGCUAUAUAGUGAAGCCACAGAUGUAUAUAGCUUCUCCUUGGUAGUCCUGCCUACUGCCUCGGA | 0.6999 |
| miR-E_03_T | UGCUGUUGACAGUGAGCGACACCAGCUUUAUGAAACUGAAUAGUGAAGCCACAGAUGUAUUCAGUUUCAUAAAGCUGGUGGUGCCUACUGCCUCGGA | 0.6931 |
| miR-E_04_T | UGCUGUUGACAGUGAGCGACACAGUGAUGCCUGUUCUUCAUAGUGAAGCCACAGAUGUAUGAAGAACAGGCAUCACUGUGGUGCCUACUGCCUCGGA | 0.6822 |
| miR-E_05_T | UGCUGUUGACAGUGAGCGAAGAAGCUAUAGACUACUUCUAUAGUGAAGCCACAGAUGUAUAGAAGUAGUCUAUAGCUUCUCUGCCUACUGCCUCGGA | 0.6689 |

| | | |
|---|---|---|
| miR-E_06_T | UGCUGUUGACAGUGAGCGAACAGCGGAUCCACCAGCUUUAUAGUGAAGCCACAGAUGUAUAAAGCUGGUGGAUCCGCUGUCUGCCUACUGCCUCGGA | 0.6679 |
| SplashRNA-designed amiRNAs | | |
| Targeting TMPRSS2 | | |
| miR-E_07_T | UGCUGUUGACAGUGAGCGCCCUGACUUAACGUUCUAUAAAUAGUGAAGCCACAGAUGUAUUUAUAGAACGUUAAGUCAGGAUGCCUACUGCCUCGGA | 1.84366 |
| miR-E_08_T | UGCUGUUGACAGUGAGCGACACCAGCUUUAUGAAACUGAAUAGUGAAGCCACAGAUGUAUUCAGUUUCAUAAAGCUGGUGGUGCCUACUGCCUCGGA | 1.751276 |
| miR-E_09_T | UGCUGUUGACAGUGAGCGACCCGUGCAUGAUUUACUCUUAUAGUGAAGCCACAGAUGUAUAAGAGUAAAUCAUGCACGGGGUGCCUACUGCCUCGGA | 1.715267 |
| miR-E_10_T | UGCUGUUGACAGUGAGCGCCAGCAGAUAUGUCUAUGACAAUAGUGAAGCCACAGAUGUAUUGUCAUAGACAUAUCUGCUGUUGCCUACUGCCUCGGA | 1.575825 |
| miR-E_11_T | UGCUGUUGACAGUGAGCGCAAGUUCAAUUGUGAAAAUGAAUAGUGAAGCCACAGAUGUAUUCAUUUUCACAAUUGAACUUUUGCCUACUGCCUCGGA | 1.566833 |
| miR-E_12_T | UGCUGUUGACAGUGAGCGACGGCAAUGUCGAUAUCUAUAAUAGUGAAGCCACAGAUGUAUUAUAGAUAUCGACAUUGCCGGUGCCUACUGCCUCGGA | 1.404016 |

* miRarchitect, shRNAI, or SplashRNA score

**Supplementary Table S4.** Individual data values and corresponding mean ± SD of the normalized Renilla/Firefly luciferase ratio from luciferase assays of miRarchitect-designed amiRNAs targeting TMPRSS2. Values represent four biological replicates with technical replicates and are normalized to the corresponding natural pri-miRNA scaffolds. The corresponding data are visualized in Supplementary Figure S3C.

| | **Biological replicate** | | | | **Mean** | **SD** |
|---|---|---|---|---|---|---|
| | **1** | **2** | **3** | **4** | | |
| pri-miR-122_nat | 1.000 | 1.000 | 1.000 | 1.000 | 1.000 | 0.000 |
| miR_122_01_T | 0.237 | 0.249 | 0.350 | 0.379 | 0.304 | 0.071 |
| miR_122_02_T | 0.200 | 0.189 | 0.140 | 0.271 | 0.200 | 0.054 |
| miR_122_03_T | 0.635 | 0.773 | 0.540 | 0.771 | 0.680 | 0.113 |
| pri-miR-155_nat | 1.000 | 1.000 | 1.000 | 1.000 | 1.000 | 0.000 |
| miR_155_01_T | 0.289 | 0.350 | 0.397 | 0.475 | 0.378 | 0.079 |
| miR_155_02_T | 0.494 | 0.424 | 0.512 | 0.551 | 0.495 | 0.053 |
| pri-miR-135b_nat | 1.000 | 1.000 | 1.000 | 1.000 | 1.000 | 0.000 |
| miR_135b_01_T | 0.268 | 0.240 | 0.210 | 0.200 | 0.229 | 0.031 |
| miR_135b_02_T | 0.234 | 0.195 | 0.220 | 0.204 | 0.213 | 0.018 |
| miR_135b_03_T | 0.450 | 0.376 | 0.375 | 0.316 | 0.379 | 0.055 |

**Supplementary Table S5.** Individual Ct values from RT-qPCR analysis of TMPRSS2, ACTIN, and/or GAPDH after cell treatment with TMPRSS2-targeting amiRNAs designed using (A) miRarchitect, (B) shRNAI, and (C) SplashRNA. Values represent three biological replicates with technical replicates. The table also reports the relative TMPRSS2 transcript expression calculated for each biological replicate. The same data are visualized in Supplementary Figures S3D and S8A.

**A**

| | **TMPRSS2** | | | **GAPDH** | | | **actin** | | **Relative TMPRSS2 expression** |
|---|---|---|---|---|---|---|---|---|---|
| **Technical replicate** | **1** | **2** | **3** | **1** | **2** | **3** | **1** | **2** | |
| amiRNA designed with miRarchitect | | | | | | | | | |
| First biological replicate | | | | | | | | | |
| pri-miR-122_nat | 16.01 | 15.64 | 15.26 | 17.88 | 17.96 | 18.25 | 23.99 | 24.57 | 1.00 |
| miR_122_01_T | 16.86 | 16.69 | 16.73 | 17.77 | 17.56 | 17.92 | 23.97 | 23.82 | 0.37 |
| miR_122_02_T | 17.65 | 17.52 | 17.69 | 18.25 | 18.08 | 17.99 | 24.66 | 24.61 | 0.29 |
| miR_122_03_T | 17.65 | 17.20 | 17.3 | 17.72 | 17.91 | 17.94 | 24.08 | 23.94 | 0.26 |

| | | | | | | | | | |
|---|---|---|---|---|---|---|---|---|---|
| pri-miR-155_nat | 16.86 | 16.56 | 16.75 | 17.88 | 17.88 | 18.08 | 24.16 | 24.05 | 1.00 |
| miR_155_01_T | 18.30 | 18.24 | 18.16 | 17.96 | 18.53 | 18.20 | 24.15 | 25.24 | 0.47 |
| miR_155_02_T | 16.92 | 17.07 | 17.28 | 17.53 | 17.83 | 18.13 | 24.03 | 23.62 | 0.68 |
| pri-miR-135b_nat | 16.24 | 15.8 | 15.82 | 18.04 | 18.26 | 17.95 | 24.81 | 24.70 | 1.00 |
| miR_135b_01_T | 17.45 | 17.29 | 17.39 | 17.82 | 17.96 | 17.89 | 23.51 | 24.22 | 0.26 |
| miR_135b_02_T | 18.45 | 18.06 | 18.22 | 17.74 | 17.91 | 18.38 | 24.25 | 24.58 | 0.18 |
| miR_135b_03_T | 16.28 | 16.56 | 16.52 | 18.09 | 18.07 | 18.33 | 24.85 | 24.69 | 0.73 |
| Control | 16.08 | 15.81 | 15.96 | 18.51 | 18.5 | 18.65 | 24.39 | 24.04 | |
| Second biological replicate | | | | | | | | | |
| pri-miR-122_nat | 16.92 | 16.45 | 16.77 | 18.74 | 18.85 | 19.07 | 24.75 | 24.57 | 1.00 |
| miR_122_01_T | 18.27 | 18.41 | 18.17 | 19.10 | 18.88 | 18.88 | 23.83 | 23.74 | 0.26 |
| miR_122_02_T | 19.43 | 18.94 | 18.84 | 19.01 | 19.04 | 18.88 | 24.24 | 24.36 | 0.18 |
| miR_122_03_T | 17.88 | 17.70 | 18.00 | 18.75 | 19.27 | 18.74 | 24.34 | 24.60 | 0.43 |
| pri-miR-155_nat | 19.08 | 19.02 | 19.08 | 20.44 | 20.18 | 20.07 | 25.02 | 24.83 | 1.00 |
| miR_155_01_T | 19.82 | 19.82 | 20.09 | 19.26 | 19.51 | 19.71 | 24.05 | 24.12 | 0.32 |
| miR_155_02_T | 18.23 | 18.44 | 18.45 | 18.52 | 18.81 | 19.09 | 24.28 | 24.01 | 0.73 |
| pri-miR-135b_nat | 18.08 | 17.96 | 17.58 | 19.06 | 18.87 | 19.42 | 23.78 | 23.59 | 1.00 |
| miR_135b_01_T | 18.99 | 18.83 | 18.93 | 19.02 | 19.10 | 19.11 | 24.54 | 24.36 | 0.60 |
| miR_135b_02_T | 19.62 | 19.32 | 19.62 | 18.94 | 19.13 | 18.89 | 23.81 | 23.55 | 0.30 |
| miR_135b_03_T | 18.77 | 18.83 | 18.74 | 19.12 | 19.11 | 19.40 | 24.35 | 24.20 | 0.66 |
| Control | 15.92 | 15.64 | 15.82 | 18.91 | 18.41 | 19.02 | 23.78 | 23.44 | |
| Third biological replicate | | | | | | | | | |
| pri-miR-122_nat | 16.13 | 16.15 | 16.07 | 17.53 | 17.2 | 17.98 | 38.20 | 38.91 | 1.00 |
| miR_122_01_T | 17.68 | 17.72 | 18.12 | 17.71 | 17.31 | 17.04 | 36.48 | 37.01 | 0.18 |
| miR_122_02_T | 18.63 | 18.22 | 18.27 | 17.40 | 17.28 | 17.79 | 37.61 | 38.08 | 0.17 |
| miR_122_03_T | 17.08 | 17.5 | 17.43 | 17.47 | 17.50 | 17.41 | 37.71 | 37.69 | 0.33 |
| pri-miR-155_nat | 18.49 | 18.98 | 18.40 | 18.34 | 18.63 | 18.54 | 38.33 | 38.95 | 1.00 |
| miR_155_01_T | 19.92 | 19.77 | 19.39 | 19.08 | 17.18 | 17.31 | 36.73 | 37.60 | 0.24 |
| miR_155_02_T | 18.21 | 18.22 | 18.12 | 17.08 | 17.63 | 17.47 | 37.28 | 37.24 | 0.56 |
| pri-miR-135b_nat | 17.28 | 17.51 | 17.57 | 17.73 | N/A | N/A | 37.18 | 38.02 | 1.00 |
| miR_135b_01_T | 18.47 | 18.31 | 18.49 | 17.89 | 17.49 | 17.20 | 37.81 | 38.19 | 0.51 |
| miR_135b_02_T | 19.04 | 18.89 | 19.38 | 17.93 | 17.58 | 18.05 | 38.31 | 38.16 | 0.39 |
| miR_135b_03_T | 19.20 | 19.14 | 18.34 | 17.81 | 17.74 | 17.58 | 38.60 | 37.99 | 0.43 |
| Control | 14.78 | 15.39 | 15.21 | 17.56 | 16.98 | 16.89 | 37.18 | 37.06 | |

**B**

| | **TMPRSS2** | | | **GAPDH** | | | **Relative TMPRSS2 expression** |
|---|---|---|---|---|---|---|---|
| **Technical replicate** | **1** | **2** | **3** | **1** | **2** | **3** | |
| amiRNA designed with shRNAI | | | | | | | |
| First biological replicate | | | | | | | |
| pri-miR-122_nat | 13.22 | 13.17 | 13.47 | 33.62 | 34.32 | 33.83 | 1.00 |
| miR-E_01_T | 13.86 | 13.89 | 13.55 | 31.37 | 31.3 | 31.63 | 0.62 |
| miR-E_02_T | 13.15 | 13.68 | 13.48 | 31.8 | 31.9 | 31.59 | 0.79 |
| miR-E_03_T | 16.33 | 17.02 | 16.17 | 33.9 | 33.28 | 33.75 | 0.11 |
| miR-E_04_T | 17.06 | 17.13 | 16.46 | 33.32 | 32.91 | 32.9 | 0.08 |
| miR-E_05_T | 14.7 | 15.03 | 14.81 | 30.66 | 30.86 | 31.03 | 0.28 |
| miR-E_06_T | 14.89 | 15.19 | 15.31 | 32.55 | 32.64 | 32.12 | 0.25 |
| Second biological replicate | | | | | | | |
| pri-miR-122_nat | 14.21 | 14.16 | 14.22 | 25.11 | 24.88 | 24.93 | 1.00 |
| miR-E_01_T | 14.91 | 15.38 | 14.62 | 25.93 | 27.29 | 27.24 | 0.66 |
| miR-E_02_T | 14.39 | 14.54 | 14.16 | 26.37 | 25.15 | 24.94 | 0.92 |
| miR-E_03_T | 17.82 | 17.78 | 17.07 | 26.23 | 27.37 | 27.8 | 0.11 |
| miR-E_04_T | 16.83 | 16.03 | 15.68 | 25.52 | 25.84 | 25.08 | 0.26 |
| miR-E_05_T | 14.58 | 14.12 | 14.39 | 26.47 | 26.6 | 25.58 | 0.97 |
| miR-E_06_T | 14.73 | 14.57 | 14.83 | 26.02 | 25.93 | 26.17 | 0.75 |
| Third biological replicate | | | | | | | |
| pri-miR-122_nat | 15.03 | 15.09 | 15.88 | 18.19 | 19.05 | 18.53 | 0.18 |

| | | | | | | | |
|---|---|---|---|---|---|---|---|
| miR-E_01_T | 15.17 | 15.48 | 15.29 | 17.64 | 17.49 | 17.46 | 0.93 |
| miR-E_02_T | 14.6 | 14.85 | 14.39 | 17.1 | 16.99 | 16.72 | 1.44 |
| miR-E_03_T | 16.23 | 16.3 | 16.22 | 16.77 | 16.54 | 16.48 | 0.45 |
| miR-E_04_T | 17.19 | 17.12 | 16.51 | 16.66 | 17.31 | 17.15 | 0.29 |
| miR-E_05_T | 15.23 | 15.23 | 15.09 | 17.47 | 17.06 | 17.07 | 0.99 |
| miR-E_06_T | 16.07 | 16.02 | 15.78 | 16.96 | 16.73 | 16.39 | 0.56 |

**C**

| | **TMPRSS2** | | | **GAPDH** | | | **Relative TMPRSS2 expression** |
|---|---|---|---|---|---|---|---|
| **Technical replicate** | **1** | **2** | **3** | **1** | **2** | **3** | |
| amiRNA designed with SplashRNA | | | | | | | |
| First biological replicate | | | | | | | |
| Pri-miR-122_nat | 15.68 | 16.93 | 15.79 | 26.82 | 26.65 | 26.96 | 1.00 |
| miR-E_07_T | 18.28 | 16.43 | 16.5 | 27.3 | 27.03 | 28.66 | 0.55 |
| miR-E_08_T | 18.52 | 18.47 | 18.37 | 28.12 | 28.05 | 27.42 | 0.21 |
| miR-E_09_T | 16.35 | 16.49 | 16.37 | 28.08 | 27.56 | 28.05 | 0.89 |
| miR-E_10_T | 18.24 | 18 | 18.4 | 27.46 | 27.55 | 27.84 | 0.25 |
| miR-E_11_T | 16.89 | 16.57 | 16.59 | 28.49 | 28.1 | 28.52 | 0.76 |
| miR-E_12_T | 18.07 | 18.15 | 17.91 | 27.39 | 27.29 | 28 | 0.28 |
| Second biological replicate | | | | | | | |
| Pri-miR-122_nat | 14.22 | 13.98 | 13.75 | 22.14 | 21.85 | 22.04 | 1.00 |
| miR-E_07_T | 15.75 | 15.7 | 15.73 | 23.3 | 23.53 | 23.59 | 0.33 |
| miR-E_08_T | 16.66 | 16.21 | 16.39 | 21.85 | 21.83 | 21.56 | 0.18 |
| miR-E_09_T | 14.9 | 14.88 | 14.72 | 22.5 | 22.09 | 22.2 | 0.57 |
| miR-E_10_T | 17.42 | 17.33 | 17.17 | 23.07 | 23.22 | 22.98 | 0.11 |
| miR-E_11_T | 15.4 | 15.65 | 15.51 | 23.27 | 23.33 | 23.75 | 0.38 |
| miR-E_12_T | 16.12 | 15.78 | 15.44 | 22.27 | 22.24 | 22.28 | 0.29 |
| Third biological replicate | | | | | | | |
| Pri-miR-122_nat | 15.28 | 15.59 | 15.19 | 22.29 | 22.19 | 22.02 | 0.18 |
| miR-E_07_T | 15.55 | 15.52 | 15.46 | 20.81 | 20.87 | 20.38 | 0.80 |
| miR-E_08_T | 17.19 | 17.15 | 16.89 | 20.87 | 20.33 | 20.33 | 0.27 |
| miR-E_09_T | 15.68 | 15.64 | 15.62 | 20.65 | 20.56 | 20.51 | 0.72 |
| miR-E_10_T | 17.52 | 17.44 | 17.29 | 20.31 | 20.89 | 20.98 | 0.21 |
| miR-E_11_T | 15.88 | 16.1 | 16.02 | 21.06 | 20.78 | 21.16 | 0.59 |
| miR-E_12_T | 16.63 | 16.44 | 16.24 | 20.65 | 19.91 | 20.14 | 0.41 |

**Supplementary Table S6.** Individual data values from Western blot analysis of relative TMPRSS2 protein expression after cell treatment with amiRNAs designed using (A) miRarchitect, (B) shRNAI, and (C) SplashRNA. Data are derived from at least three biological replicates, each measured in three technical replicates. The same data are visualized in Supplementary Figures S3E, S4, and S8B.

**A**

| **Biological replicate** | **1** | | | | **2** | | | | **3** | | | | **4** | | | |
|---|---|---|---|---|---|---|---|---|---|---|---|---|---|---|---|---|
| **Technical replicate** | **1** | **2** | **3** | **Mean** | **1** | **2** | **3** | **Mean** | **1** | **2** | **3** | **Mean** | **1** | **2** | **3** | **Mean** |
| pri-miR-122_nat | 1.000 | 1.000 | 1.000 | 1.000 | 1.000 | 1.000 | 1.000 | 1.000 | 1.000 | 1.000 | 1.000 | 1.000 | 1.000 | 1.000 | 1.000 | 1.000 |
| miR_122_01_T | 0.042 | 0.037 | 0.037 | 0.039 | 0.063 | 0.055 | 0.033 | 0.050 | 0.065 | 0.058 | 0.032 | 0.051 | 0.092 | 0.059 | 0.075 | 0.075 |
| miR_122_02_T | 0.062 | 0.048 | 0.048 | 0.053 | 0.054 | 0.035 | 0.033 | 0.040 | 0.050 | 0.043 | 0.012 | 0.035 | 0.071 | 0.055 | 0.054 | 0.060 |
| miR_122_03_T | 0.374 | 0.514 | 0.276 | 0.388 | 0.296 | 0.165 | 0.157 | 0.206 | 0.127 | 0.187 | 0.071 | 0.128 | 0.145 | 0.104 | 0.101 | 0.117 |
| pri-miR-155_nat | 1.000 | 1.000 | 1.000 | 1.000 | 1.000 | 1.000 | 1.000 | 1.000 | 1.000 | 1.000 | 1.000 | 1.000 | 1.000 | 1.000 | 1.000 | 1.000 |
| miR_155_01_T | 0.050 | 0.051 | 0.056 | 0.053 | 0.084 | 0.121 | 0.199 | 0.135 | 0.365 | 0.223 | 0.156 | 0.248 | 0.353 | 0.167 | 0.421 | 0.314 |
| miR_155_02_T | 0.111 | 0.231 | 0.205 | 0.182 | 0.280 | 0.281 | 0.521 | 0.361 | 0.676 | 0.457 | 0.493 | 0.542 | 0.557 | 0.612 | 0.885 | 0.685 |
| pri-miR-135b_nat | 1.000 | 1.000 | 1.000 | 1.000 | 1.000 | 1.000 | 1.000 | 1.000 | 1.000 | 1.000 | 1.000 | 1.000 | 1.000 | 1.000 | 1.000 | 1.000 |
| miR_135b_01_T | 0.112 | 0.375 | 0.045 | 0.177 | 0.136 | 0.149 | 0.361 | 0.215 | 0.134 | 0.085 | 0.106 | 0.108 | 0.153 | 0.152 | 0.156 | 0.153 |
| miR_135b_02_T | 0.131 | 0.075 | 0.066 | 0.091 | 0.156 | 0.181 | 0.202 | 0.180 | 0.103 | 0.046 | 0.127 | 0.092 | 0.114 | 0.080 | 0.131 | 0.108 |
| miR_135b_03_T | 0.577 | 0.796 | 0.468 | 0.614 | 0.653 | 0.854 | - | 0.753 | 0.329 | 0.147 | 0.198 | 0.225 | 0.316 | 0.323 | 0.320 | 0.320 |

**B**

| **Biological replicate** | **1** | | | | **2** | | | | **3** | | | |
|---|---|---|---|---|---|---|---|---|---|---|---|---|
| **Technical replicate** | **1** | **2** | **3** | **Mean** | **1** | **2** | **3** | **Mean** | **1** | **2** | **3** | **Mean** |
| pri-miR-122_nat | 1.000 | 1.000 | 1.000 | 1.000 | 1.000 | 1.000 | 1.000 | 1.000 | 1.000 | 1.000 | 1.000 | 1.000 |
| miR-E_01_T | 1.083 | 0.726 | 0.753 | 0.854 | 1.338 | 0.848 | 0.857 | 1.014 | 0.938 | 0.769 | 0.553 | 0.753 |
| miR-E_02_T | 1.428 | 0.915 | 0.734 | 1.026 | 1.227 | 1.263 | 1.509 | 1.333 | 1.061 | 0.778 | 0.561 | 0.800 |
| miR-E_03_T | 0.136 | 0.261 | 0.202 | 0.200 | 0.033 | 0.023 | 0.298 | 0.118 | 0.078 | 0.234 | 0.118 | 0.143 |
| miR-E_04_T | 0.084 | 0.106 | 0.124 | 0.105 | 0.022 | 0.027 | 0.124 | 0.058 | 0.032 | 0.198 | 0.166 | 0.132 |
| miR-E_05_T | 0.211 | 0.162 | 0.173 | 0.182 | 0.868 | 0.908 | 1.028 | 0.935 | 0.751 | 1.104 | 0.878 | 0.911 |
| miR-E_06_T | 0.181 | 0.253 | 0.193 | 0.209 | 1.135 | 0.902 | 0.722 | 0.920 | 0.138 | 0.510 | 0.194 | 0.281 |

**C**

| **Biological replicate** | **1** | | | | **2** | | | | **3** | | | |
|---|---|---|---|---|---|---|---|---|---|---|---|---|
| **Technical replicate** | **1** | **2** | **3** | **Mean** | **1** | **2** | **3** | **Mean** | **1** | **2** | **3** | **Mean** |
| Pri-miR-122_nat | 1.000 | 1.000 | 1.000 | 1.000 | 1.000 | 1.000 | 1.000 | 1.000 | 1.000 | 1.000 | 1.000 | 1.000 |
| miR-E_07_T | 0.763 | 0.756 | 0.737 | 0.752 | 0.928 | 0.407 | 0.306 | 0.547 | 1.015 | 0.426 | 0.361 | 0.601 |

| | | | | | | | | | | | |
|---|---|---|---|---|---|---|---|---|---|---|---|
| miR-E_08_T | 0.069 | 0.044 | 0.018 | 0.044 | 0.150 | 0.082 | 0.096 | 0.109 | 0.154 | 0.131 | 0.089 | 0.125 |
| miR-E_09_T | 0.902 | 0.845 | 0.411 | 0.719 | 1.206 | 1.002 | 0.922 | 1.043 | 0.951 | 0.545 | 0.526 | 0.674 |
| miR-E_10_T | 0.126 | 0.066 | 0.023 | 0.072 | 0.084 | 0.108 | 0.135 | 0.109 | 0.125 | 0.570 | 0.073 | 0.256 |
| miR-E_11_T | 1.120 | 1.192 | 0.701 | 1.004 | | 1.315 | 1.290 | 1.303 | 0.890 | 0.651 | 0.842 | 0.794 |
| miR-E_12_T | 0.203 | 0.227 | 0.061 | 0.163 | 0.098 | 0.092 | 0.217 | 0.136 | 0.141 | 0.203 | 0.219 | 0.188 |

**Supplementary Table S7.** Individual Ct values from RT-qPCR analysis of ACE-2 and GAPDH after cell treatment with ACE-2–targeting amiRNAs designed using miRarchitect. Values represent four biological replicates with technical replicates. The table also reports the relative ACE-2 transcript expression calculated for each biological replicate. The same data are visualized in Supplementary Figure S5B.

| | **ACE-2** | | | **GAPDH** | | | **Relative ACE-2 expression** |
|---|---|---|---|---|---|---|---|
| **Technical replicate** | **1** | **2** | **3** | **1** | **2** | **3** | |
| First biological replicate | | | | | | | |
| Pri-miR-122_nat | 15.21 | 15.21 | 15.74 | 18 | 18.32 | 18.1 | 1.00 |
| miR_122_01_A | 16.37 | 16.8 | 16.6 | 18.45 | 18.38 | 18.42 | 0.44 |
| Pri-miR-155_nat | 15.07 | 15.31 | 15.45 | 18.34 | 18.32 | 18.1 | 1.00 |
| miR_155_01_A | 17.41 | 17.43 | 17.2 | 18.82 | 18.75 | 18.92 | 0.25 |
| Second biological replicate | | | | | | | |
| Pri-miR-122_nat | 33.77 | 33.11 | 32.24 | 20.46 | 20.26 | 20.42 | 1.00 |
| miR_122_01_A | 33.89 | 36.42 | 33.74 | 21.26 | 21.09 | 21.42 | 0.34 |
| Pri-miR-155_nat | 34.22 | 32.26 | 32.1 | 20.8 | 21.04 | 21.12 | 1.00 |
| miR_155_01_A | 34.17 | 34.73 | 38.57 | 21.75 | 21.7 | 21.81 | 0.14 |
| Third biological replicate | | | | | | | |
| Pri-miR-122_nat | 32.21 | 31.3 | 32.23 | 19.05 | 18.96 | 19.13 | 1.00 |
| miR_122_01_A | 34.02 | 33.6 | 33.46 | 19.57 | 19.94 | 20.19 | 0.31 |
| Pri-miR-155_nat | 32.7 | 31.48 | 31.64 | 19.28 | 18.9 | 19.3 | 1.00 |
| miR_155_01_A | 32.58 | 33.09 | 33.07 | 18.51 | 18.81 | 19.69 | 0.50 |
| Fourth biological replicate | | | | | | | |
| Pri-miR-122_nat | 32.49 | 31.16 | 31.56 | 17.39 | 17.37 | 17.53 | 1.00 |
| miR_122_01_A | 33.89 | 33.46 | 32.78 | 18.33 | 19.02 | 19.13 | 0.36 |
| Pri-miR-155_nat | 31.96 | 31.4 | 31.5 | 17.1 | 18.01 | 17.51 | 1.00 |
| miR_155_01_A | 32.52 | 32.2 | 32.31 | 18.31 | 18.24 | 18.19 | 0.64 |

**Supplementary Table S8.** Individual data values from Western blot analysis of relative ACE-2 protein expression after cell treatment with ACE-2–targeting amiRNAs designed using miRarchitect. Values are derived from at least four biological replicates, each measured in three technical replicates. The same data are visualized in Supplementary Figure S5B.

| **Biological replicate** | **1** | | | | **2** | | | | **3** | | | | | **4** | | | | | |
|---|---|---|---|---|---|---|---|---|---|---|---|---|---|---|---|---|---|---|---|
| **Technical replicate** | **1** | **2** | **3** | **Mean** | **1** | **2** | **3** | **Mean** | **1** | **2** | **3** | **4** | **Mean** | **1** | **2** | **3** | **4** | **5** | **Mean** |
| Pri-miR-122_nat | 1.000 | 1.000 | 1.000 | 1.000 | 1.000 | 1.000 | 1.000 | 1.000 | 1.000 | 1.000 | 1.000 | 1.000 | 1.000 | 1.000 | 1.000 | 1.000 | 1.000 | 1.000 | 1.000 |
| miR_122_01_A | 0.353 | 0.399 | 0.367 | 0.373 | 0.579 | 0.491 | 0.405 | 0.492 | 0.583 | 0.447 | 0.557 | 0.623 | 0.552 | 0.936 | 0.539 | 0.713 | 0.460 | 0.860 | 0.702 |
| Pri-miR-155_nat | 1.000 | 1.000 | 1.000 | 1.000 | 1.000 | 1.000 | 1.000 | 1.000 | 1.000 | 1.000 | 1.000 | 1.000 | 1.000 | 1.000 | 1.000 | 1.000 | 1.000 | 1.000 | 1.000 |
| miR_155_01_A | 0.590 | 0.301 | 0.360 | 0.417 | 0.520 | 0.432 | 0.618 | 0.523 | 0.425 | 0.466 | 0.424 | 0.410 | 0.431 | 0.573 | 0.964 | 0.844 | 0.738 | 0.545 | 0.733 |

**Supplementary Table S9.** Percentage of guide strand reads among total reads for the most abundant endogenous miRNAs – those contributing more than 0.1% of all small RNA-seq reads in HEK293T cells.

| amiRNA | Guide strand (%) |
|---|---|
| miR_122_01_T | 0.02 |
| miR_122_02_T | 0.55 |
| miR_122_03_T | 0.01 |
| miR_155_01_T | 0.84 |
| miR_155_02_T | 0.55 |
| miR_135b_01_T | 0.02 |
| miR_135b_02_T | 0.23 |
| miR_135b_03_T | 0.02 |
| miR_122_01_A | 4.68 |
| miR_155_01_A | 32.26 |

**Supplementary Table S10.** Key features of web-based tools for amiRNA design.

| **Name** | **Organism** | **Scaffold** | **Ranking of amiRNAs with scores** | **Off-target list** | **Experimentally validated amiRNA** |
|---|---|---|---|---|---|
| miRarchitect | *Homo sapiens* | hsa-mir-21, hsa-mir-30a, hsa-mir-122, hsa-mir-135b*, hsa-mir-155 | + | + | + |
| shRNAI | *Homo sapiens, Mus musculus* | miR-E** | + | - | + |
| SplashRNA | *Homo sapiens, Mus musculus* | miR-E** | + | - | + |
| miRNA designer | *Arabidopsis thaliana, Homo sapiens* | ath-MIR168, ath-MIR158 | - | - | + |
| WMD3. Web MicroRNA Designer | *Arabidopsis thaliana, Oryza sativa* | ath-MIR319a, osa-MIR528 | + | - | + |
| amiRNA Design Helper | *Marchantia polymorpha* | mp-MIR160 | + | + | + |

* The pri-miR-135b scaffold. which may produce mature miRNAs from both arms, is not selected by default, as such dual-arm processing is not recommended for therapeutic applications.

** miR-30-based optimized backbone with a conserved element at the 3'end of the basal stem, required for optimal amiRNA processing.

**Supplementary Table S11.** Oligonucleotides used in the study.

| Gene | Primer orientation | Sequence (5'-3') | Description |
|---|---|---|---|
| TMPRSS2_XhoI_NotI | F | CGATCTCGAGCCATGCCCCCTGCCCCG CCCGGA | Primers for TMPRSS2v1 amplification before cloning into psiCHECK2 plasmid. F and R primers provide the sequences for XhoI and NotI cleavage sites, respectively. |
| | R | CCATGCGGCCGCTTAGCCGTCTGCCCT CATTTG | |
| TMPRSS2v1 | F | CACTGTGCATCACCTTGACC | RT-qPCR primers |
| | R | ACACACCGATTCTCGTCCTC | |
| GAPDH | F | GGTCGGAGTCAACGGATTTG | RT-qPCR primers |
| | R | GGATCTCGCTCCTGGAAGAT | |
| ACTIN | F | TGAGAGGGAAATCGTGCGTG | RT-qPCR primers |
| | R | TGCTTGCTGATCCACATCTGC | |
| ACE-2 | F | CAGGGAACAGGTAGAGGACATT | RT-qPCR primers |
| | R | CAGAGGGTGAACATACAGTTGG | |

**Supplementary Table S12.** Antibodies used in the study.

| Protein | Primary antibody | Secondary antibody |
|---|---|---|
| TMPRSS2-V5 Tag | Anti-V5 Tag mouse mAb, conjugated with HRP, ThermoFisher (R961-25), 1:2500 in 3% BSA PBS-T | n/a |
| ACE-2 | Anti-ACE-2 recombinant rabbit mAb, ThermoFisher (MA5-32307, 1:1000 in 3% BSA PBS-T | Peroxidase AffiniPure® Donkey Anti-Rabbit IgG (H+L), Jackson ImmunoReserch, Cat#711-035-152 |
| GAPDH | Anti-GAPDH mouse mAb, ThermoFisher (MA1-16757), 1:1000 in 3% BSA PBS-T | Goat anti-mouse IgG (H+L) Ab, conjugated with HRP, ThermoFisher, Cat # 31430 |

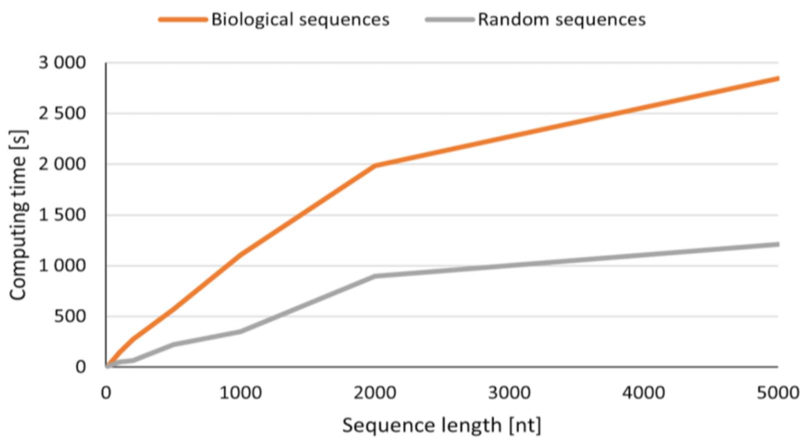


**Supplementary Figure S1.** Scalability of miRarchitect for large-scale inputs. The corresponding raw data are provided in Supplementary Table S1.

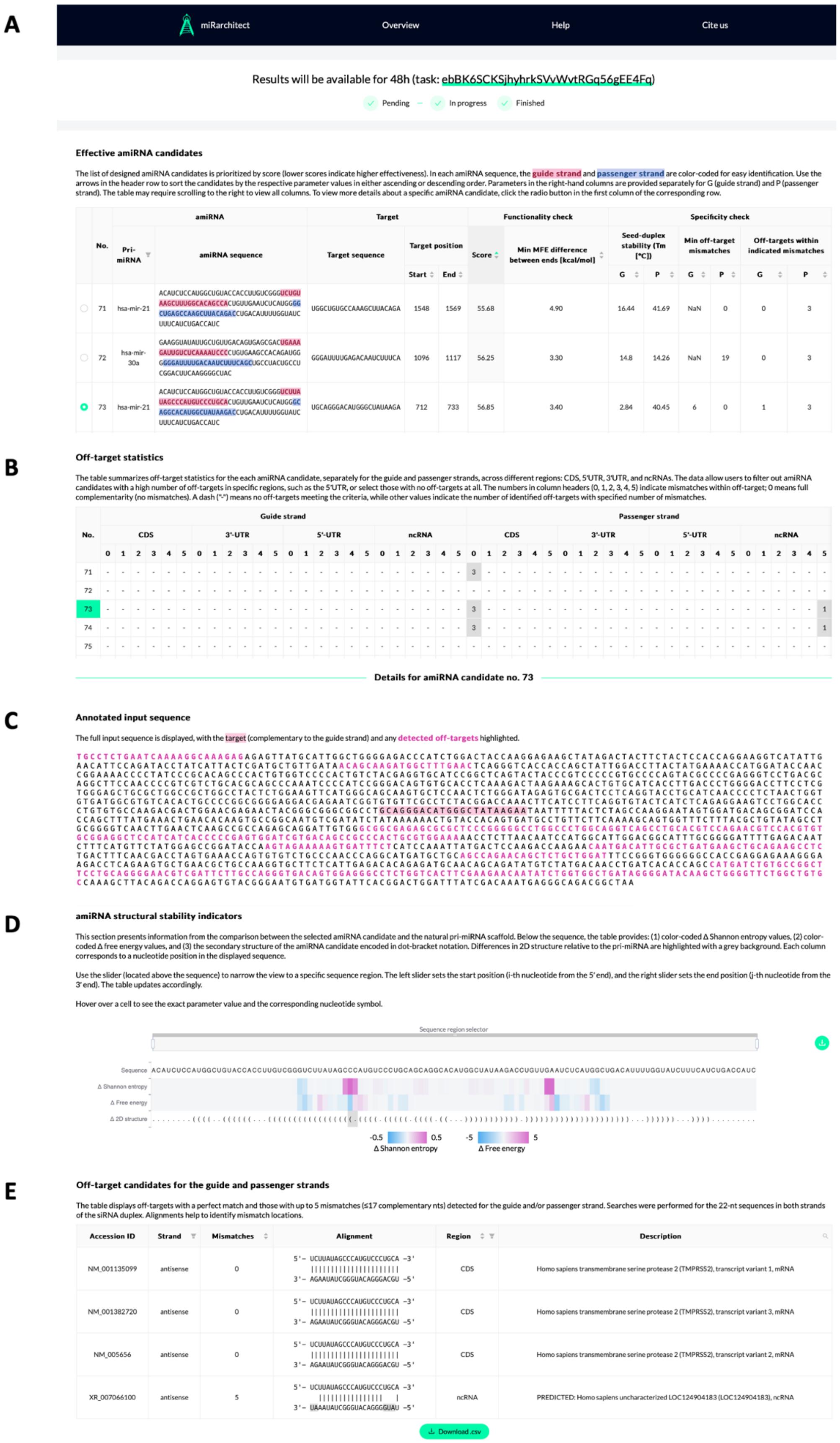


**Supplementary Figure S2.** miRarchitect output: (A) ranked list of TMPRSS2-targeting amiRNAs, (B) off-target statistics, (C) predicted target regions, (D) structural stability indicators, and (E) list of off-targets for guide and passenger strands.

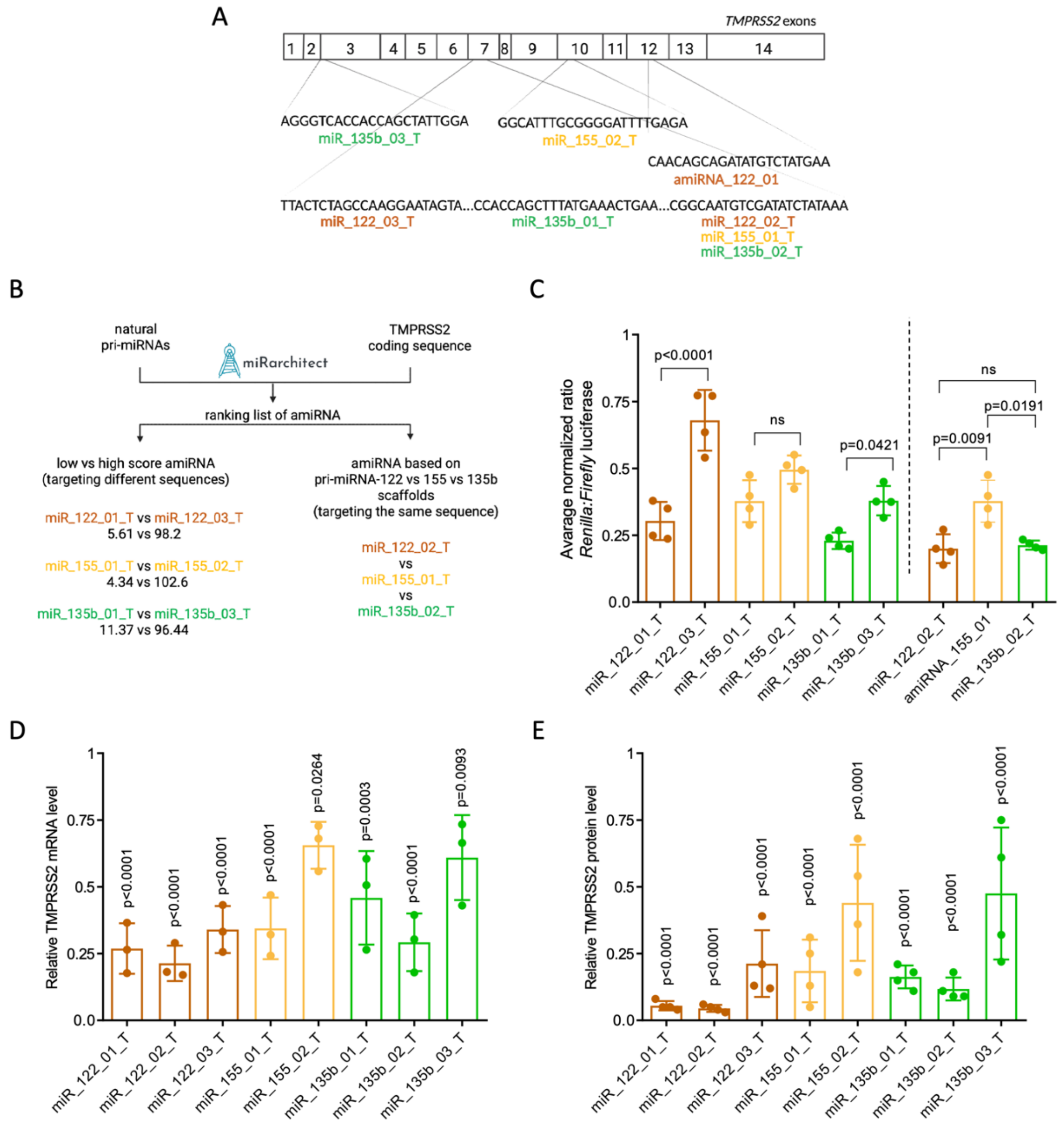


**Supplementary Figure S3.** Silencing efficiency of TMPRSS2-targeting amiRNAs designed using miRarchitect. (A) Schematic representation of TMPRSS2 exons highlighting target sequences for amiRNAs. (B) Comparison of amiRNAs selected from the miRarchitect ranking list under two conditions: (i) low-scoring vs. high-scoring amiRNAs sharing the same pri-miRNA scaffold, and (ii) amiRNAs targeting the same sequence but derived from different scaffolds. (C) Luc reporter assay showing knockdown efficiency of anti-TMPRSS2 amiRNAs. HEK293T cells were co-transfected with the Luc reporter encoding TMPRSS2v.1 cDNA and either amiRNA-expressing plasmids or control plasmids. Silencing efficiency is presented as the *Renilla*/firefly luciferase ratio. normalized to the natural pri-mRNA control and analyzed using one-way ANOVA with Tukey's multiple comparison test. (D) RT-qPCR analysis of TMPRSS2 transcript levels. (E) Western blot analysis of TMPRSS2 protein levels in HEK293T cells treated with plasmids encoding TMPRSS2v.1 cDNA and amiRNA plasmids. Protein band intensities were normalized to GAPDH and analysed using a one-sample t-test. Data are shown as mean ± SD (n = 4). P-values are indicated; "ns" denotes non-significant (p-value>0.05). Corresponding raw data are provided in Supplementary Tables S4–S6.

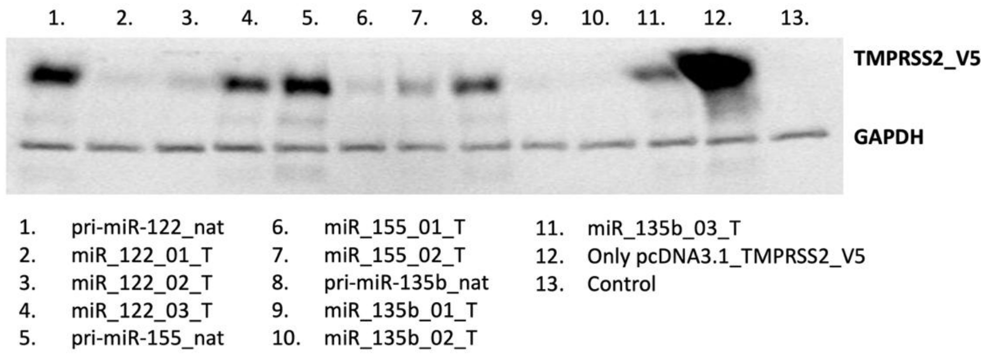


**Supplementary Figure S4.** Representative Western blot showing TMPRSS2_V5 and GAPDH protein expression in HEK293T cells 24 hours after transfection with pcDNA3.1_TMPRSS2_V5 and amiRNA vectors. GAPDH served as a loading control.

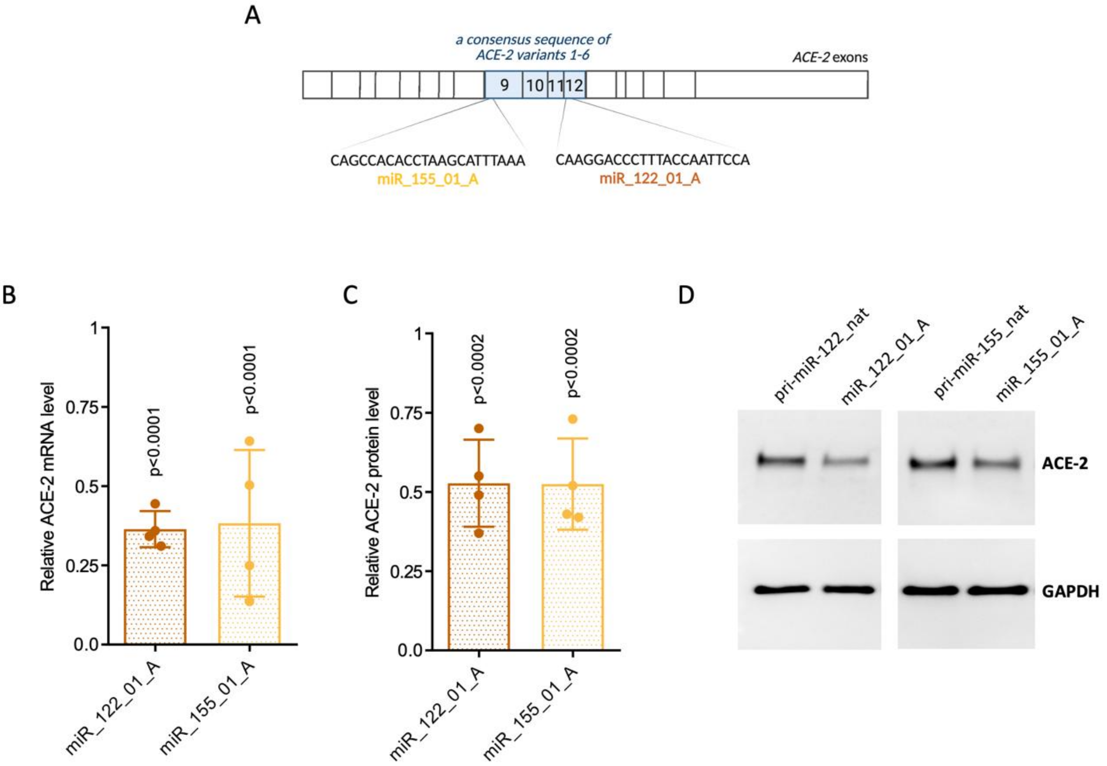


**Supplementary Figure S5.** Silencing efficiency of ACE-2-targeting amiRNAs designed using miRarchitect. (A) Schematic representation of ACE-2 exons highlighting target sequences for amiRNAs. (B) RT-qPCR analysis of ACE-2 transcript levels. (C) Western blot analysis of ACE-2 protein levels in HEK293T cells treated with plasmids encoding ACE-2 and amiRNA plasmids. Protein band intensities were normalized to GAPDH and analysed using a one-sample t-test. Data are shown as mean ± SD (n = 4). Corresponding raw data are provided in Supplementary Tables S7–S8. (D) Representative Western blot showing ACE-2 and GAPDH protein expression in HEK293T cells 24 hours after transfection with pcDNA3.1_ACE-2 and amiRNA vectors. GAPDH served as a loading control.

**A**

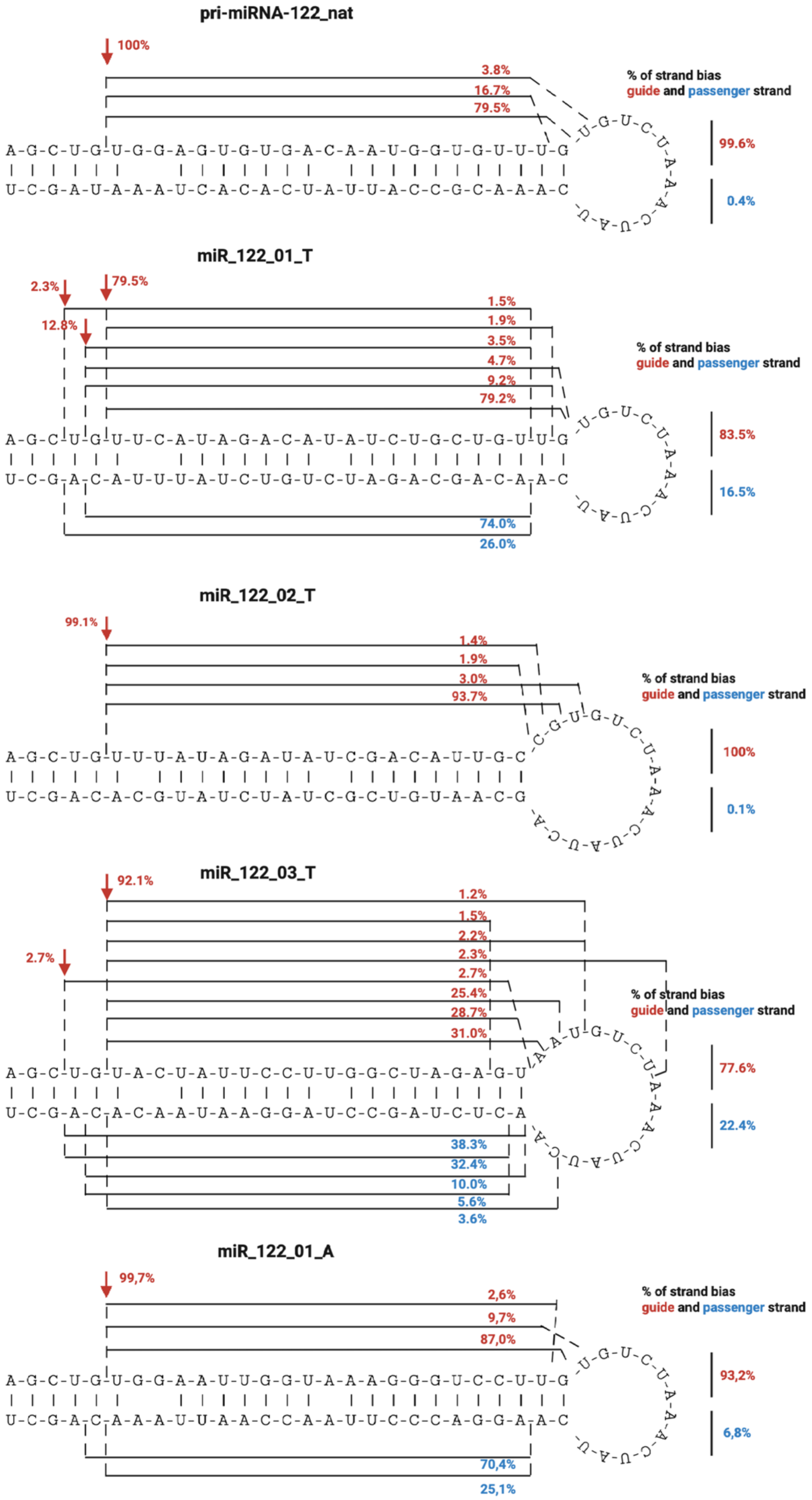
pri-miRNA-122_nat
100%
3.8%
16.7%
79.5%
% of strand bias
guide and passenger strand
99.6%
0.4%
miR_122_01_T
2.3%
79.5%
12.8%
1.5%
1.9%
3.5%
4.7%
9.2%
79.2%
% of strand bias
guide and passenger strand
83.5%
16.5%
74.0%
26.0%
miR_122_02_T
99.1%
1.4%
1.9%
3.0%
93.7%
% of strand bias
guide and passenger strand
100%
0.1%
miR_122_03_T
92.1%
1.2%
1.5%
2.2%
2.3%
2.7%
2.7%
25.4%
28.7%
31.0%
% of strand bias
guide and passenger strand
77.6%
22.4%
38.3%
32.4%
10.0%
5.6%
3.6%
miR_122_01_A
99,7%
2,6%
9,7%
87,0%
% of strand bias
guide and passenger strand
93,2%
6,8%
70,4%
25,1%

B

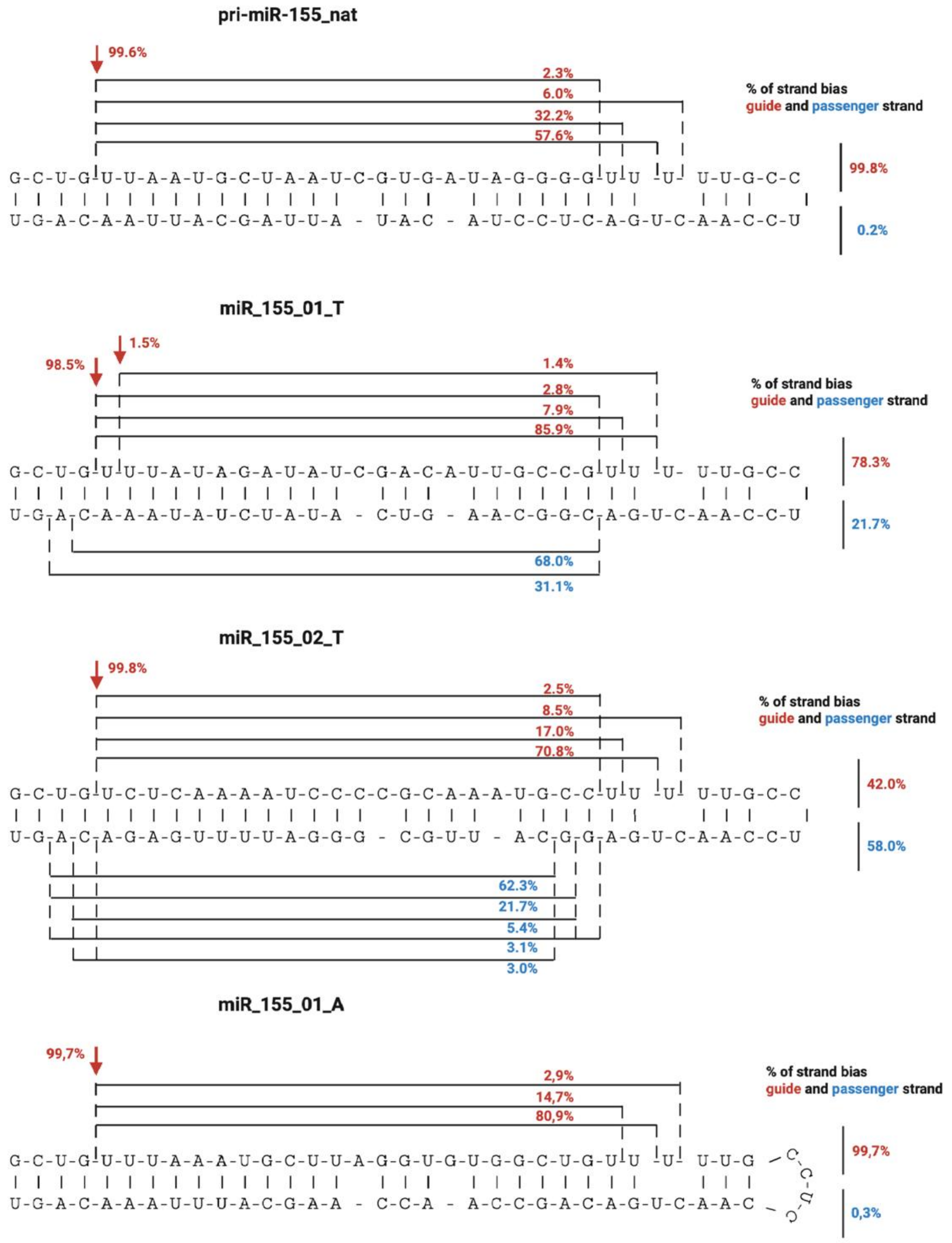
pri-miR-155_nat
99.6%
2.3%
6.0%
32.2%
57.6%
% of strand bias
guide and passenger strand
G-C-U-G-U-U-A-A-U-G-C-U-A-A-U-C-G-U-G-A-U-A-G-G-G-G-U-U -U- U-U-G-C-C
U-G-A-C-A-A-U-U-A-C-G-A-U-U-A - U-A-C - A-U-C-C-U-C-A-G-U-C-A-A-C-C-U
99.8%
0.2%
miR_155_01_T
1.5%
98.5%
1.4%
2.8%
7.9%
85.9%
% of strand bias
guide and passenger strand
G-C-U-G-U-U-U-A-U-A-G-A-U-A-U-C-G-A-C-A-U-U-G-C-C-G-U-U -U- U-U-G-C-C
U-G-A-C-A-A-A-U-A-U-C-U-A-U-A - C-U-G - A-A-C-G-G-C-A-G-U-C-A-A-C-C-U
78.3%
21.7%
68.0%
31.1%
miR_155_02_T
99.8%
2.5%
8.5%
17.0%
70.8%
% of strand bias
guide and passenger strand
G-C-U-G-U-C-U-C-A-A-A-U-C-C-C-C-G-C-A-A-U-G-C-C-U-U -U- U-U-G-C-C
U-G-A-C-A-G-A-G-U-U-U-U-A-G-G-G - C-G-U-U - A-C-G-G-A-G-U-C-A-A-C-C-U
42.0%
58.0%
62.3%
21.7%
5.4%
3.1%
3.0%
miR_155_01_A
99,7%
2,9%
14,7%
80,9%
% of strand bias
guide and passenger strand
G-C-U-G-U-U-U-A-A-A-U-G-C-U-U-A-G-G-U-G-U-G-G-C-U-G-U-U -U- U-U-G
C-C-U-C
U-G-A-C-A-A-A-U-U-U-A-C-G-A-A - C-C-A - A-C-C-G-A-C-A-G-U-C-A-A-C
99,7%
0,3%

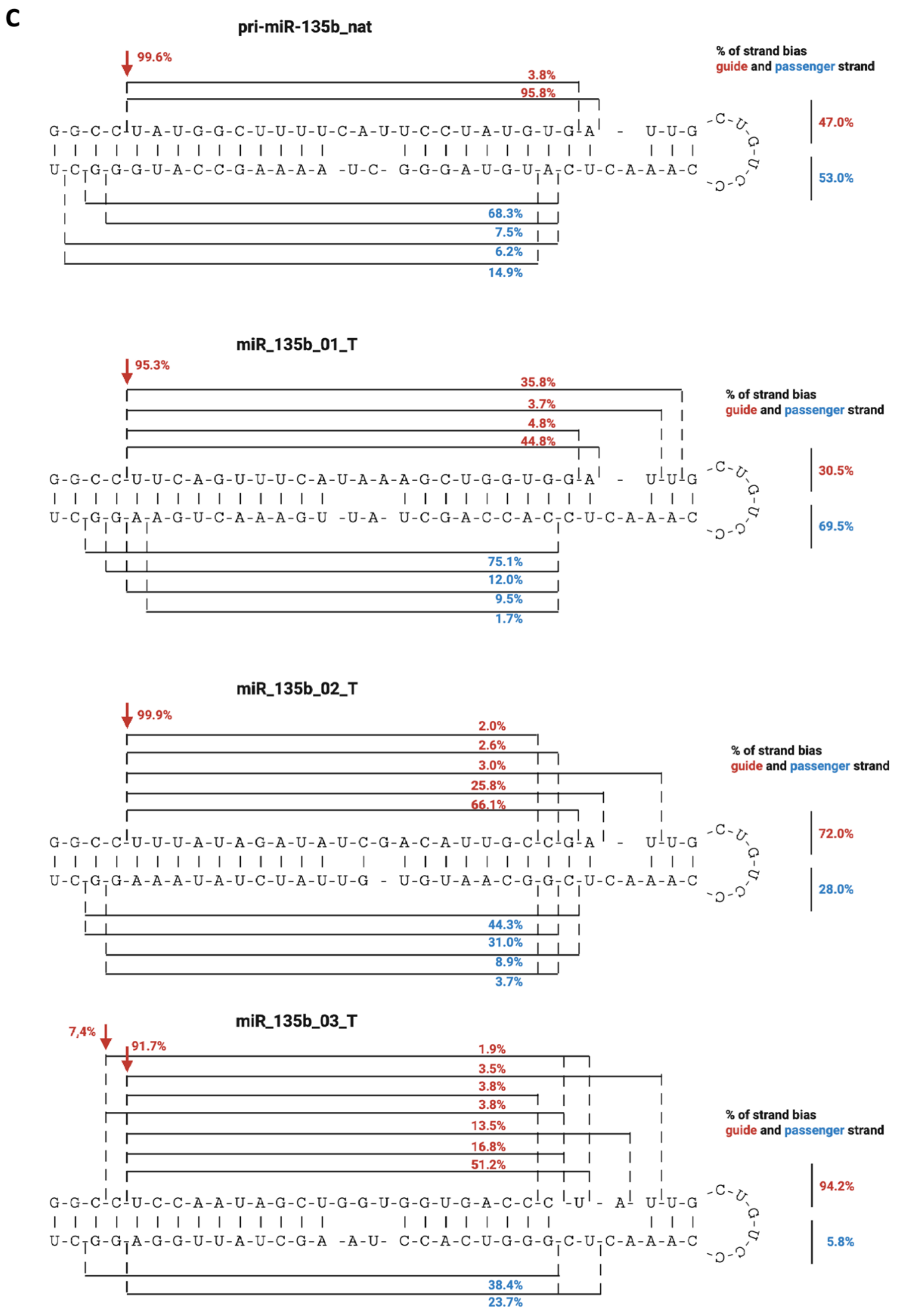


**Supplementary Figure S6.** Processing of miRarchitect-designed amiRNA targeting TMPRSS2 and ACE-2 in HEK293T cells. Next-generation sequencing analysis of the processing of amiRNAs based on (A) pri-miR-122, (B) pri-miR-135b, and (C) pri-miR-155 scaffolds, along with the corresponding natural pri-miRNAs used as controls. Mature miRNAs derived from the guide and passenger strands are shown in red and blue, respectively, with their relative abundances indicated. Red arrows denote the Drosha cleavage sites.

**A**

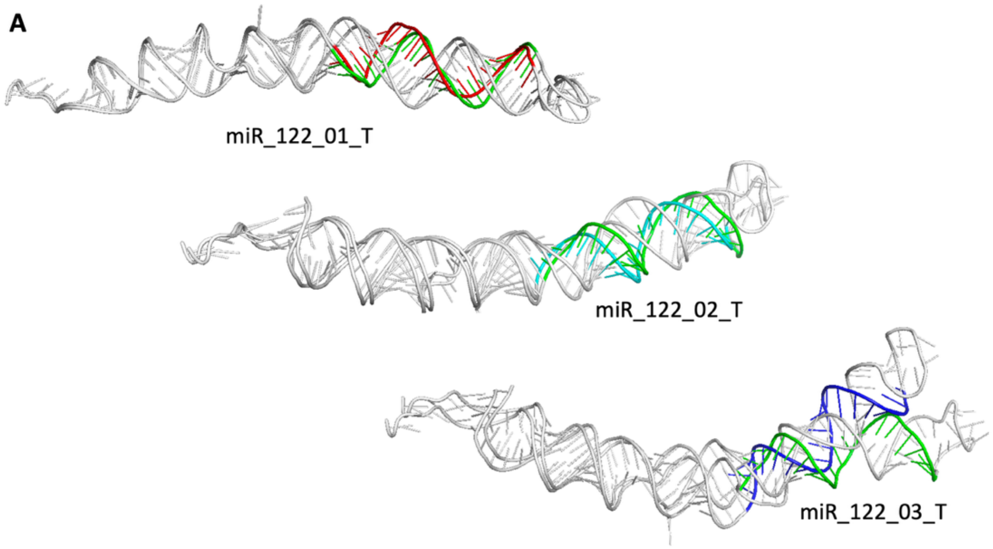


**B**

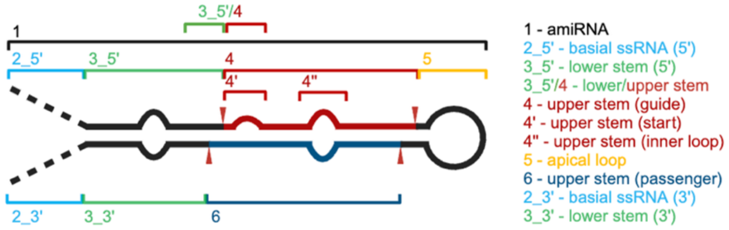


**C**

| | RMSD [Å] (amiRNA vs pri-miRNA scaffold) | | | | | | | | | | |
|---|---|---|---|---|---|---|---|---|---|---|---|
| | amiRNA | Basal ssRNA (5') | Lower stem (5') | Lower/Upper stem | Upper stem (guide) | Upper stem (start) | Upper stem (inner loop) | Apical loop | Upper stem (passenger) | Lower stem 3' | Basal ssRNA 3' |
| miR_122_01_T | 2,854 | 0,155 | 0,233 | 0,964 | 1,914 | 1,252 | 0,322 | 0,211 | 2,031 | 0,165 | 0,159 |
| miR_122_02_T | 7,033 | 0,177 | 0,267 | 1,359 | 3,311 | 1,285 | 0,283 | 4,866 | 1,897 | 0,141 | 0,15 |
| miR_122_03_T | 11,851 | 0,127 | 0,258 | 0,513 | 4,382 | 0,783 | nd | 4,894 | 5,248 | 0,108 | 0,119 |
| miR_155_01_T | 4,598 | 0,11 | 0,17 | 0,088 | 3,485 | 0,077 | 2,167 | 0,27 | 2,042 | 0,143 | 0,092 |
| miR_155_02_T | 4,052 | 0,105 | 0,139 | 0,07 | 6,549 | 0,042 | 7,671 | 0,31 | 2,751 | 0,085 | 0,076 |
| miR_135b_01_T | 7,409 | 0,232 | 0,23 | 0,392 | 5,106 | 0,123 | 1,264 | 0,543 | 4,461 | 0,12 | 0,159 |
| miR_135b_02_T | 2,559 | 0,19 | 0,19 | 0,37 | 2,521 | 0,403 | nd | 0,385 | 3,591 | 0,225 | 0,185 |
| miR_135b_03_T | 5,025 | 0,208 | 0,257 | 0,41 | 5,066 | 0,103 | 2,186 | 3,671 | 2,818 | 0,235 | 0,188 |

Color scale: 0 – 14/nd

**Supplementary Figure S7.** amiRNA structures aligned with their scaffolds. (A) The 3D structures of individual pri-miR-122-based amiRNAs aligned with the 3D structure of their natural scaffold (pri-miR-122). Guide strands are color-coded: hsa-miR-122 (green), miR_122_01_T (red), miR_122_02_T (cyan), and miR_122_03_T (blue). (B) Structural deviations between the amiRNAs and their scaffold are quantified using Root Mean Square Deviation (RMSD). (C) RMSD values and mismatched regions are visualized as a heat map, with greater discrepancies highlighted in red.

A

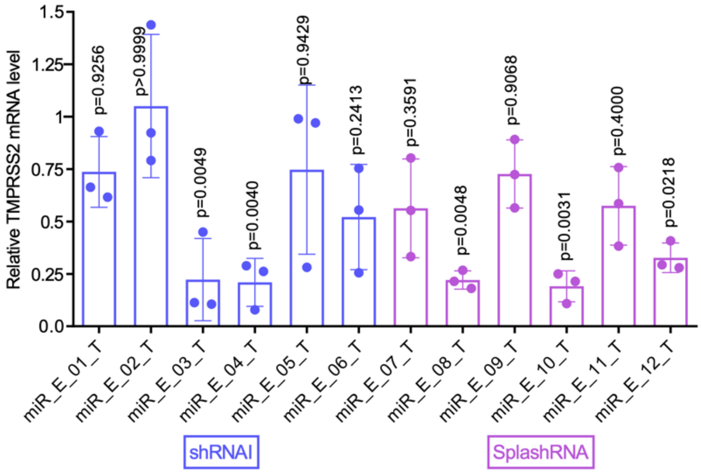


B

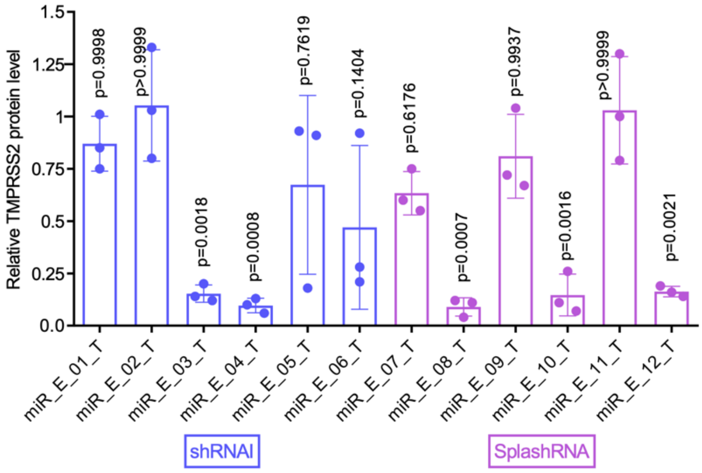


**Supplementary Figure S8.** Silencing efficiency of TMPRSS2-targeting amiRNAs designed using shRNAI and SplashRNA. (A) RT-qPCR analysis of TMPRSS2 transcript levels, (B) Western blot analysis of TMPRSS2 protein levels in HEK293T cells treated with plasmids encoding TMPRSS2v.1 cDNA and amiRNA plasmids. Protein band intensities were normalized to GAPDH and analysed using a one-sample t-test. Data are shown as mean ± SD (n = 3). Corresponding raw data are provided in Supplementary Tables S5–S6.
